# Therapy-Induced Clonal Selection as a Driver of Response to JAK Inhibitors in Myelofibrosis

**DOI:** 10.64898/2026.06.20.733514

**Authors:** Sebastiano Rontauroli, Chiara Carretta, Matteo Bertesi, Sandra Parenti, Daniela Benati, Monica Maccaferri, Tommaso Ferrari, Marica Malerba, Anita Neroni, Elisa Papa, Ruggiero Norfo, Margherita Mirabile, Lara Tavernari, Camilla Tombari, Paola Guglielmelli, Alessandra Recchia, Leonardo Potenza, Rossana Maffei, Enrico Tagliafico, Mario Luppi, Alessandro Maria Vannucchi, Rossella Manfredini

## Abstract

Myelofibrosis (MF) originates from the stepwise acquisition of somatic mutations in Hematopoietic Stem and Progenitor Cells (HSPCs). Alongside *driver* events triggering JAK-STAT pathway hyperactivation, several additional mutations, usually affecting the epigenetic machinery, contribute defining therapeutic response. Specifically, JAK-inhibition (JAKi) relieves MF symptoms but rarely eradicates the neoplastic clone.

To elucidate clonal dynamics associated with JAKi, we conducted a longitudinal single-cell proteogenomic study on 6 responders and 6 non-responders MF patients. Mutational analysis revealed that the mutation acquisition order determines JAKi sensitivity. Indeed, *driver*-only clones are highly sensitive to JAKi, while co-mutated clones persist after treatment. JAKi response is mainly limited to the differentiated myeloid compartment, while mutant HSPCs are often maintained in JAKi-responders. Co-mutated clones may evade JAKi and outcompete other neoplastic cell populations, thus contributing to disease persistence.

## Introduction

Myeloproliferative Neoplasms (MPNs) are a family of clonal hematological disorders originating from Hematopoietic Stem Cells (HSCs). Within this group, Myelofibrosis (MF) is characterized by the poorest prognosis, progressive bone marrow fibrosis, splenomegaly and development of extra-medullary hematopoiesis. The MPN genetic framework is shaped by recurrent pathogenic mutations acquired by HSCs. Mutually exclusive somatic “driver mutations” affecting *JAK2*, *CALR* or *MPL* hyperactivate JAK-STAT signaling pathway, and result in the accumulation of differentiated myeloid cells. MF display the more complex genetic landscape among MPNs due to the accumulation of somatic variants, copy number variations and aneuploidies^1^. Patient-specific mutational assets play a role in defining disease course and therapeutic response^2–4^. Tet Methylcytosine Dioxygenase 2 (*TET2*) and ASXL Transcriptional Regulator 1 (*ASXL1*) loss of function mutations are the most frequent non-driver events in MF, detected in 17% and 13% of patients respectively^5^. These variants play a pivotal role in MF as they influence cell fate, response to treatment and disease evolution but underlying molecular mechanisms remain not fully understood^3^.

Acquired somatic mutations in genes such as *ASXL1*, *EZH2*, *IDH1/2*, *SRSF2* and *U2AF1* are classified as high-molecular risk gene (HMR) variants since they are associated with adverse prognosis^4,6^ and are independent risk factors for survival in MF^7,8^. The order of mutation acquisition impact disease onset and progression because of their effect on hematopoietic stem and progenitor cells (HSPCs) biology. *TET2* or *DNMT3A* mutations confer a clonal advantage to HSPCs differently from *JAK2* variants, which most likely impact differentiation. As a result, mutation acquisition order impacts the balance between myeloid and megakaryocyte-erythroid progenitors as well as disease phenotype^2,9^. Moreover, in MF patients evolved to secondary acute myeloid leukemia the MPN driver mutation is more frequently preceded by the acquisition of an epigenetic variant^10,11^.

To date, allogeneic hematopoietic stem cells’ transplantation remains the only curative therapeutic option for MF. However, most patients are managed with targeted therapies, primarily JAK-inhibitors (JAKi) including the first in class Ruxolitinib (Ruxo). Ruxo reduces splenomegaly and improve quality of life by reducing symptoms^12,13^ but almost half of the patients discontinue it by the third year of treatment^14^. As of today, Ruxo treatment failure might refer to: therapy resistance or inadequate response due to lack of spleen volume reduction; relapse or loss of response; disease progression including AML transformation; intolerance due to the development of cytopenias, including anemia or thrombocytopenia^14–17^. The molecular determinants of treatment response include JAK2V617F variant allele frequency (VAF) >50% which is associated with the achievement of spleen volume reduction^18,19^. Conversely, a reduced clinical response to JAKi is associated with the number and type of non-driver mutations^20–23^.

JAK inhibitors have a limited impact on tumor burden and consequently fail to prevent disease progression or leukemic transformation. Studies have demonstrated that molecular response after JAKi treatment is poor, being significant in a small proportion of MF patients and after long term treatment^14^. In the present work, we employed single-cell (SC) multiomic to study molecular response in JAKi treated MF patients and dissect cellular and genetic determinants of resistance. Specifically, we demonstrate that the prior acquisition of mutations affecting epigenetic remodelers impairs response to JAKi, which predominantly target clones carrying the sole driver mutation. Therefore, co-mutated clones can evade this treatment and outcompete other neoplastic cell populations, thereby promoting clonal persistence and disease progression.

## Results

### Circulating CD34+ cells are reduced after JAK inhibition in Responder patients

To investigate clonal dynamics associated with JAKi response we analyzed a cohort composed of 12 myelofibrosis (MF) patients treated with Ruxolitinib (Ruxo) or Fedratinib (Fed). According to response criteria^24^, in 6 patients JAK inhibition was able to improve constitutional symptoms or reduce spleen volume, therefore these patients were classified as Responders (Resp) (Table S1). The remaining patients, Non-Responders (Non-Resp), did not experience any significant benefit from Ruxo treatment, or required treatment interruption due to side effects. Among them, patient #4 underwent leukemic transformation while patient #1 suspended Ruxo treatment and moved to Fed due to increased circulating CD34+ cells and white blood cells (Table S1). These patient groups do not significantly differ in terms of treatment duration (mean time on treatment: Resp= 20.3 months, range 8-38 months; Non-Resp= 17.6 months, range 5-25 months).

According to bulk genomic analysis of whole blood samples collected before Ruxo initiation (Fig. 1A), all patients displayed at least one MPN driver variant, which was accompanied by additional somatic variants in 10/12 cases. As expected, JAK2-only mutated patients belonged to Resp while in Non-Resp the number of mutated genes per patient was generally higher. Indeed, a higher number of non-driver mutations is inversely correlated with the achievement of spleen volume response and predicts earlier Ruxo discontinuation^21^ *TET2* and *ASXL1* variants were the most frequent non-driver mutations in both Resp and Non-Resp (Fig. 1B). To study the clinical and molecular effects of JAK inhibition, two longitudinal samples per patient were analyzed: one collected before JAKi start, the second collected after at least 6 months of treatment. Blood counts were analyzed at the same time points and revealed that the frequency of circulating CD34+ cells was reduced in Resp after treatment (Fig. 1C) in line with previous studies which correlated this parameter with spleen volume response and suggested its use as prognostic marker in JAKi treated MF patients^25,26^. Additionally, driver mutation VAF in granulocytes displays a non-significant trend towards reduction only in Resp as determined using droplet digital PCR (ddPCR) (Fig. 1D).

**Figure 1:**
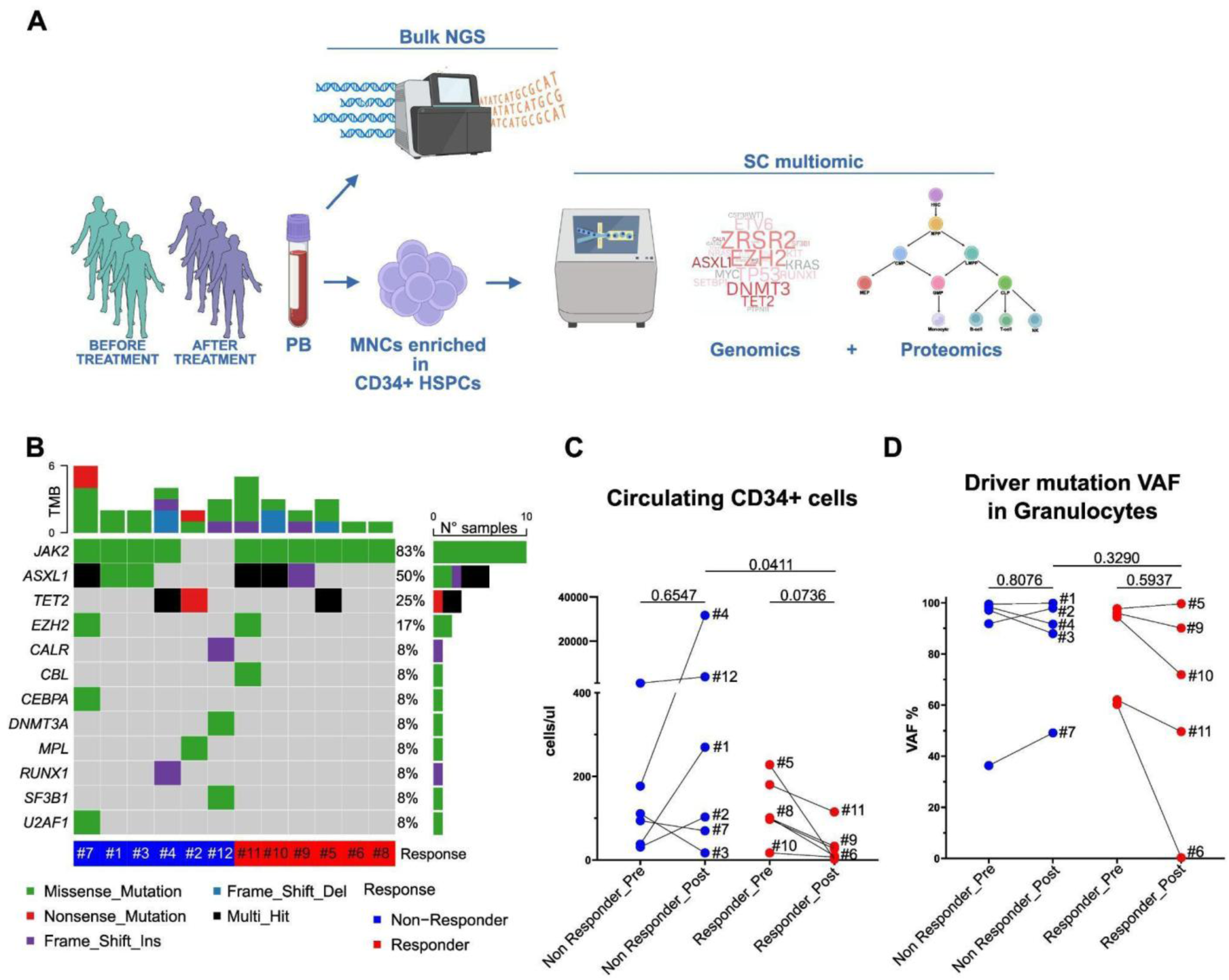
Circulating CD34+ cells are reduced in responder patients after JAK inhibitor treatment. **A** Experimental design. 12 patients diagnosed with myelofibrosis (MF) and treated with JAK inhibitors were included in the study. From each patient a peripheral blood sample was collected at two different time points, before and after JAK inhibitor treatment, as shown in Supplementary Figure 1 and Supplementary Tables 1. A fraction of pre-treatment whole blood sample was subjected to bulk next generation sequencing (NGS) to identify acquired somatic mutations. Mononuclear cells (MNCs) were isolated from both pre-treatment and post-treatment samples and cryopreserved. The day of single cell (SC) multiomic analysis, after thawing, CD34+ hematopoietic stem and progenitor cells (HSPCs) were purified then mixed in a 1:1 ratio with corresponding unfractionated MNC cells. The resulting cell suspension was subjected to SC genomic and proteomic analysis using Tapestri platform according to producer instructions. **B** Onco plot showing the type and the number of driver and non-driver mutations identified in patient samples according to bulk NGS. Patients were classified as Responders and Non-Responders according to spleen volume reduction, symptoms amelioration and disease progression (Supplementary Tables 1). **C** Variation of circulating CD34+ cell number in patients before (Pre) and after (Post) JAK inhibitor treatment according to blood count evaluation. Each pair of dots represents a patient (n=6 Responders; n=6 Non-Responders). **D** Variation of driver mutations variant allele frequency (VAF) in granulocytes from Responders and Non-Responders before (Pre) and after (Post) JAK inhibitor treatment. Each pair of dots represents a patient (n=5 Responders; n=5 Non-Responders). Comparisons were performed using Friedman test followed by uncorrected Dunn’s test for paired samples, and Mann-Whitney test for unpaired samples. P-values are reported. Abbreviations: HSPCs: hematopoietic stem and progenitor cells; MNCs: mononuclear cells; NGS: next generation sequencing; PB: peripheral blood; SC: single cell; TMB: tumour mutation burden; VAF: variant allele frequency.

### Ruxo treatment affects the frequency of circulating mononuclear cell subpopulations

To study clonal dynamics associated with JAKi treatment we performed combined SC genomic and proteomic analysis of circulating CD34⁺ HSPCs and peripheral blood mononuclear cells (PBMCs) collected before and after JAKi treatment Based on SC proteomic information we classified cells and identified 14 clusters including: differentiated mononuclear cells (monocytes, CD4+ and CD8+ T cells, natural killer (NK) cells and B cells), erythroid and myeloid CD34^-^ precursors, and 6 clusters of CD34+ HSPCs (HSCs, multipotent progenitors (MPP), lymphoid primed multipotent progenitors (LMPP), common myeloid progenitors (CMP) and megakaryocyte and erythroid progenitors (MEP)) (Fig. 2A, B, Figure S1A). Since the number of analyzed cells was highly variable among samples (Table S2) we sub-sampled our data cohort by including 1000 cells per sample while maintaining the frequency of cell clusters within each sample (Figure S1B, C).

**Figure 2:**
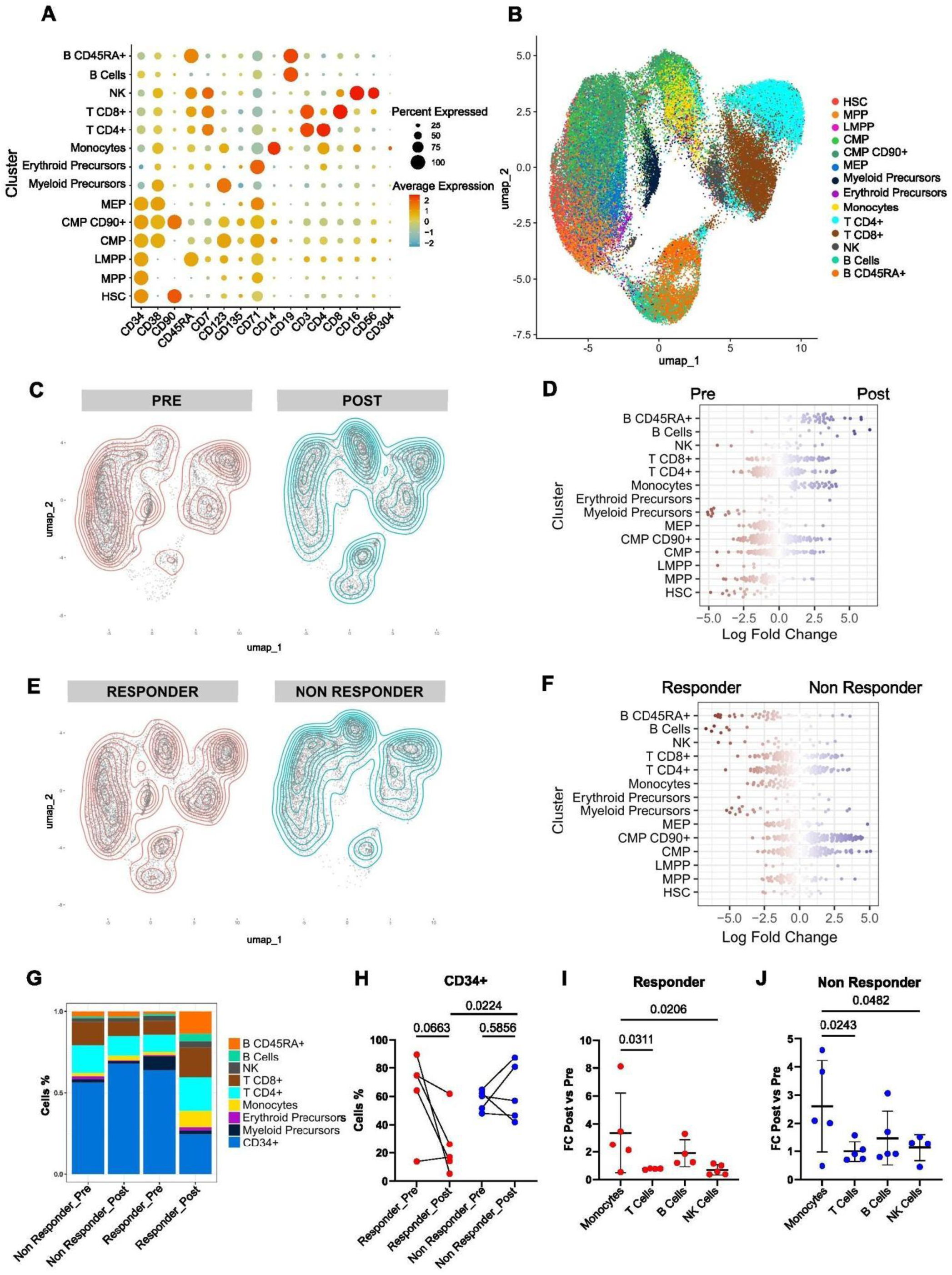
Circulating mononuclear cells classified according to SC proteomics. **A** Single cell (SC) proteo-genomic analysis was performed in 9 patients (5 Responders and 4 Non-Responders). The bubble chart displays the expression of selected antigens in cell clusters identified using the FlowSOM R package. **B** Uniform manifold approximation and projection (UMAP) representation of analysed cells (n= 59163) distributed according to surface antigen expression and coloured according to cell clusters identified using FlowSOM R package. **C** 2D density plot showing the distribution of cells in the UMAP before (Pre) and after (Post) JAK inhibitor treatment. **D** Beeswarm plot of differential cell abundance represented as log Fold Change (FC) in neighborhoods identified using MILO R package and classified according to FlowSOM cell clusters. The differential distribution of cells before (Pre) and after (Post) JAK inhibitor treatment was evaluated. **E** 2D density plot showing the distribution of cells in the UMAP in Responders and Non-Responders. **F** Beeswarm plot of differential cell abundance represented as log Fold Change in neighborhoods identified using MILO R package and classified according to FlowSOM cell clusters. The differential distribution of cells from Responders and Non-Responders was evaluated. **G** Cells were distinguished according to timepoint and treatment response in four groups. The bar plot displays the frequency of cell types within each group. Hematopoietic stem and progenitor cell (HSPC) clusters were grouped together as CD34+ cells. **H** Scatter plot showing the variation of CD34+ cells frequency in samples subjected to single cell (SC) genomic analysis. Each pair of dots represents a patient (n=5 Responders; n=5 Non-Responders). **I and J** Scatter plot represents the variation of differentiated cells frequency in Responder (**I**)(n=5) and Non-Responder (**J**)(n=5) patients. The frequency of each cell population was determined within each patient sample based on SC analysis. The variation of the frequency of each cell population after JAK inhibitor treatment was expressed as FC and shown in the scatter plot. Each dot represents a patient. Comparisons were performed using Friedman test followed by uncorrected Dunn’s test for paired samples, and Mann-Whitney test for unpaired samples. P-values less than 0.05 are considered significant. Abbreviations: B CD45RA+: B Cells expressing CD45RA marker; CMP: common myeloid progenitors; CMP CD90+: CMP positive for CD90; FC: fold change; HSC: hematopoietic stem cells; LMPP: Lymphoid primed multipotent progenitors; MEP: megakaryocyte erythroid progenitors; MPP: multipotent progenitors; NK: natural killers; UMAP: Uniform manifold approximation and projection.

As regards cell distribution among clusters we observed that differentiated cell clusters were enriched in post treatment samples, particularly from Resp, while CD34+ HSPC clusters were enriched in pre-treatment samples from Non-Resp (Fig. 2C-F). SC proteomic results confirmed blood count results showing a significant reduction in the frequency of circulating CD34+ cells in Resp samples after JAK inhibition (Fig. 2G, H). SC proteomic analysis revealed that in both Resp and Non-Resp CD14+ monocytes frequency was significantly increased after JAKi treatment as compared to other differentiated cell populations (Fig. 2I, J).

### Clonal hierarchy and dynamic distinguish Resp from Non-Resp

SC multiomic analysis allowed the reconstruction of the mutation acquisition order and clonal architecture in immunophenotypically defined hematopoietic cell populations. According to SC genomics, mutation acquisition order efficiently distinguished Resp from Non-Resp. As for Resp, 2 out of 6 harbored JAK2V617F mutation alone, while the remaining acquired at least one additional event in non-driver genes that are sub-clonal of the MPN driver variant. Conversely, Non-Resp most likely acquired the MPN driver variant after a preceding hit in an epigenetic modifier gene, particularly *TET2* or *ASXL1*. An exception is represented by Non-Resp patient #3, in whom only JAK2V617F mutation was detected by SC genomics due to the reduced genotyping efficiency of the amplicon covering the known *ASXL1* mutation harbored by this patient (Fig. 3A, B).

**Figure 3:**
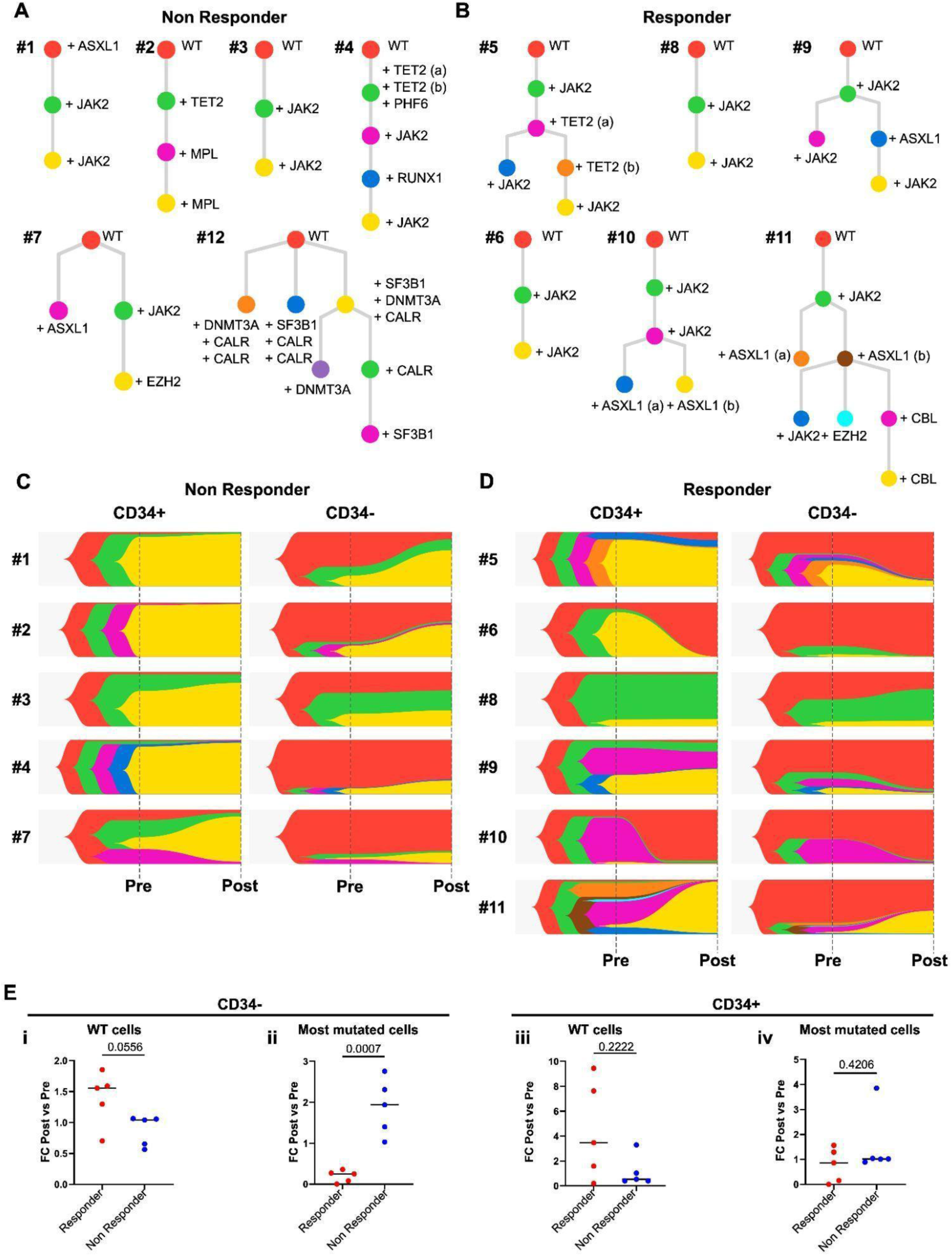
Epigenetic variants precede the acquisition of driver mutations in Non-Responder patients. **A and B** Phylogenetic trees represent the mutation acquisition order and clonal evolution in Responder and Non-Responder patients reconstructed according to single cell (SC) genomic information. The mutational event leading to the appearance of a new clone is indicated by the symbol of the affected gene. When, in a patient, a gene is affected by more than one variant, each of them is distinguished by a lowercase letter in brackets; the corresponding mutation is reported in Supplementary Table 1. **C and D** Fish plot representing the clonal dynamics over time in Responder and Non-Responder patients. CD34+ hematopoietic stem and progenitor cells (HSPCs) and CD34- differentiated cells were distinguished based on SC proteomic information. Clone colors are the same used in phylogenetic trees. **E** In each patient we analysed the variation of the frequency of the wild type (WT) clone (**i** and **iii**) and the most mutated one (**ii** and **iv**). Their frequency variation was expressed as the fold change in the comparison between post-treatment and pre-treatment sample. Scatter plots represent the FC of clones frequency in both differentiated CD34- cells (**i** and **ii**) and CD34+ HSPCs (**iii** and **iv**). Each dot represents a patient (n=5 Responders and n=5 Non-responders). Comparisons were performed using Kruskal-Wallis test. P-values less than 0.05 are considered significant and reported in panels. Abbreviations: FC: fold change; WT: wild type.

In Non-Resp, the most mutated malignant clone which dominated hematopoiesis before treatment further expanded or at least remained stable during time in both CD34+ and CD34-cells. On the contrary, in Resp, non-mutated cells expanded over time in 4 out of 6 patients (Fig. 3C, D). Of note, among Resp, patient #8 harbored the JAK2V617F mutation alone and fish plots revealed the expansion of the heterozygous clone within CD34- cell compartment (Fig. 3D). As previously described^11,27^, SC genomics revealed chromosome 9p duplication in this patient involving *JAK2* locus and being acquired in two distinct clonal branches (Fig. S2A). Clonal dynamics reconstruction revealed that, in both CD34+ and CD34- cell compartments, the clone harboring two JAK2 mutated copies out of three expanded during time despite Ruxo treatment (Fig. S2B).

Clonal prevalence variations were more pronounced in CD34- differentiated cell compartment. Statistical analysis confirmed most mutated clones’ expansion in Non-Resp and contraction in Resp. Within CD34+ cells, differences between Resp and Non-Resp were statistically non-significant (Fig. 3E).

### Genomic Complexity Is Associated with Progressive Cellular Immaturity

Given the complexity and heterogeneity of clonal composition among different patients we sought to simplify our dataset by grouping genetic clones according to shared molecular features. We classified cells based on the acquisition order of MPN driver mutations and epigenetic variants and their zygosity. We identified 8 shared genetic clones, namely: unmutated cells (WT), cells that acquired one mutation only in epigenetic regulator (Epi), cells harboring heterozygous (Driver_Het) or homozygous (Driver_Hom) MPN driver mutation, cells that acquired one mutation in epigenetic regulators before heterozygous (Epi+Driver_Het) or homozygous (Epi+Driver_Hom) MPN driver mutation, cells that acquired one mutation in epigenetic regulator after heterozygous (Driver_Het+Epi) or homozygous (Driver_Hom+Epi) MPN driver mutation (Fig. 4A).

**Figure 4:**
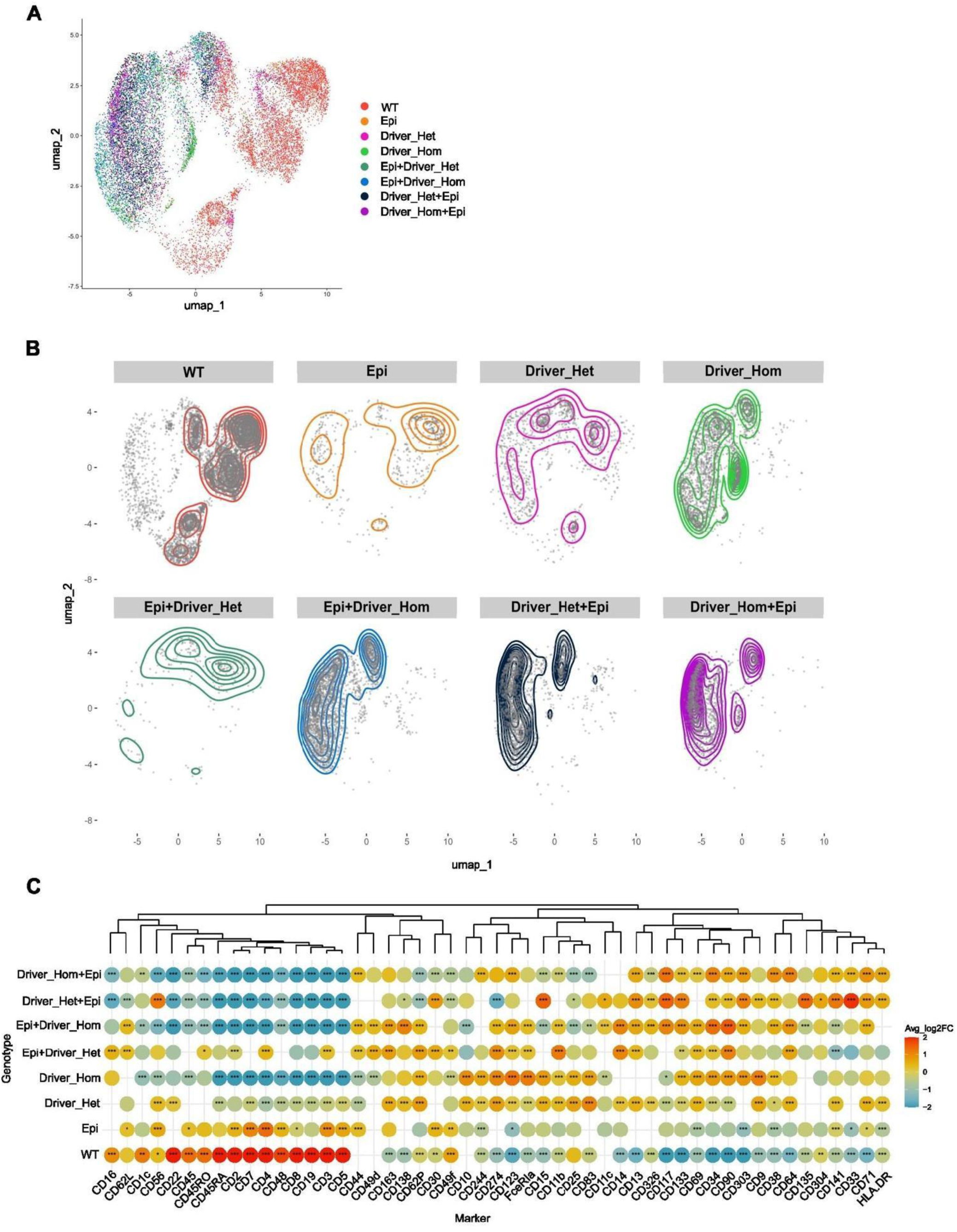
Genomic classification of clones according to SC multiomic analysis. **A** Cells were classified in 8 genetic groups according to the order of mutation acquisition and the zygosity of MPN driver mutation. The UMAP displays the distribution of these genetic groups within the proteomic defined cell landscape. **B** Density plots highlight the distribution of genetic groups within the UMAP. **C** The bubble chart displays protein expression fold change (FC) highlighting the correlation between genomic classification and protein expression. Genotype-associated protein markers were identified through FindMarkers function of Seurat package (Wilcoxon Rank Sum test, logfc.threshold = 0.25). *, P < 0,05; **, P < 0,01; and ***, P < 0,001 Abbreviations: Driver_Het: clone harboring heterozygous MPN driver mutation; Driver_Hom: clone harboring homozygous MPN driver mutation; Driver_Het+Epi: clone harboring heterozygous MPN driver mutation followed by variant(s) in epigenetic regulator(s); Driver_Hom+Epi: clone harboring homozygous MPN driver mutation followed by variant(s) in epigenetic regulator(s); Epi: clone harboring mutation(s) in epigenetic regulator(s); Epi+Driver_Het: clone harboring variant(s) in epigenetic regulator(s) followed by heterozygous MPN driver mutation; Epi+Driver_Hom: clone harboring variant(s) in epigenetic regulator(s) followed by homozygous MPN driver mutation; WT: wild type clone.

By leveraging SC multiomic analysis, we integrated genomic classification with proteomic information, correlating cellular mutational landscapes with surface protein expression in cells from pre-treatment samples to avoid any effect driven by JAKi treatment. As reported in Figure 4B the different genetic clones were unevenly distributed within UMAP. As expected, most WT cells are localized within non-myeloid differentiated cell clusters, including T-, B- and NK-cells. Epi or Epi+Driver_Het cells were similarly enriched in differentiated clusters, including monocytes. Accordingly, WT and Epi clones were associated with the expression of NK and lymphoid cell markers including CD16, CD56, CD3, CD4, CD8, and CD19 (Fig. 4C and Fig. S3). Conversely, the presence of an MPN driver mutation correlated with a myeloid and immature cellular phenotype. Expression of myeloid markers, such as CD163, CD15, CD11b, CD14, CD123, and CD244, correlated with Driver_Het and Driver_Hom clones (Fig. 4C and Fig. S3). The combination of epigenetic variant and a preceding MPN driver mutation, either heterozygous and homozygous, correlated with the expression of hematopoietic stem and progenitor cells markers including CD34, CD90, CD133, CD117, CD38, CD135 and CD71 (Fig. 4C and Fig. S3). Comparison of cells harboring heterozygous versus homozygous MPN driver mutations revealed that higher zygosity correlated with assignment to the HSPC compartment, both in single mutated and co-mutated Epi+Driver_Het and Epi+Driver_Hom ones (Fig. 4B). In contrast, in Driver_Het+Epi and Driver_Hom+Epi this difference was mainly absent (Fig. 4B).

Collectively, our results reveal a shift toward myeloid lineage in the presence of MPN driver mutations together with increased genomic complexity marking progressive cellular immaturity.

### MPN driver homozygous clones are the most sensitive to JAK inhibition

Once we have identified shared genetic clones, our next step was to characterize how they respond to JAK inhibition. To this end we evaluated the frequency of genetic clones at different timepoints in Resp and Non-Resp. Interestingly, we observed the expansion of WT clones accompanied by a contraction of both Driver_Hom and Driver_Hom+Epi clones among Resp cells. Conversely, in Non-Resp cells, despite JAK inhibition, co-mutated clones (Epi+Driver_Hom and Driver_Het+Epi) expanded at the expense of WT and single mutated ones (Fig. 5A). By analyzing Resp and Non-Resp patients separately we confirmed the significant increase in the frequency of WT cells, the reduction of Driver_Hom cells in Resp and the significant expansion of cells displaying additional epigenetic modifier gene mutations in Non-Resp after treatment (Fig. 5B, C).

**Figure 5:**
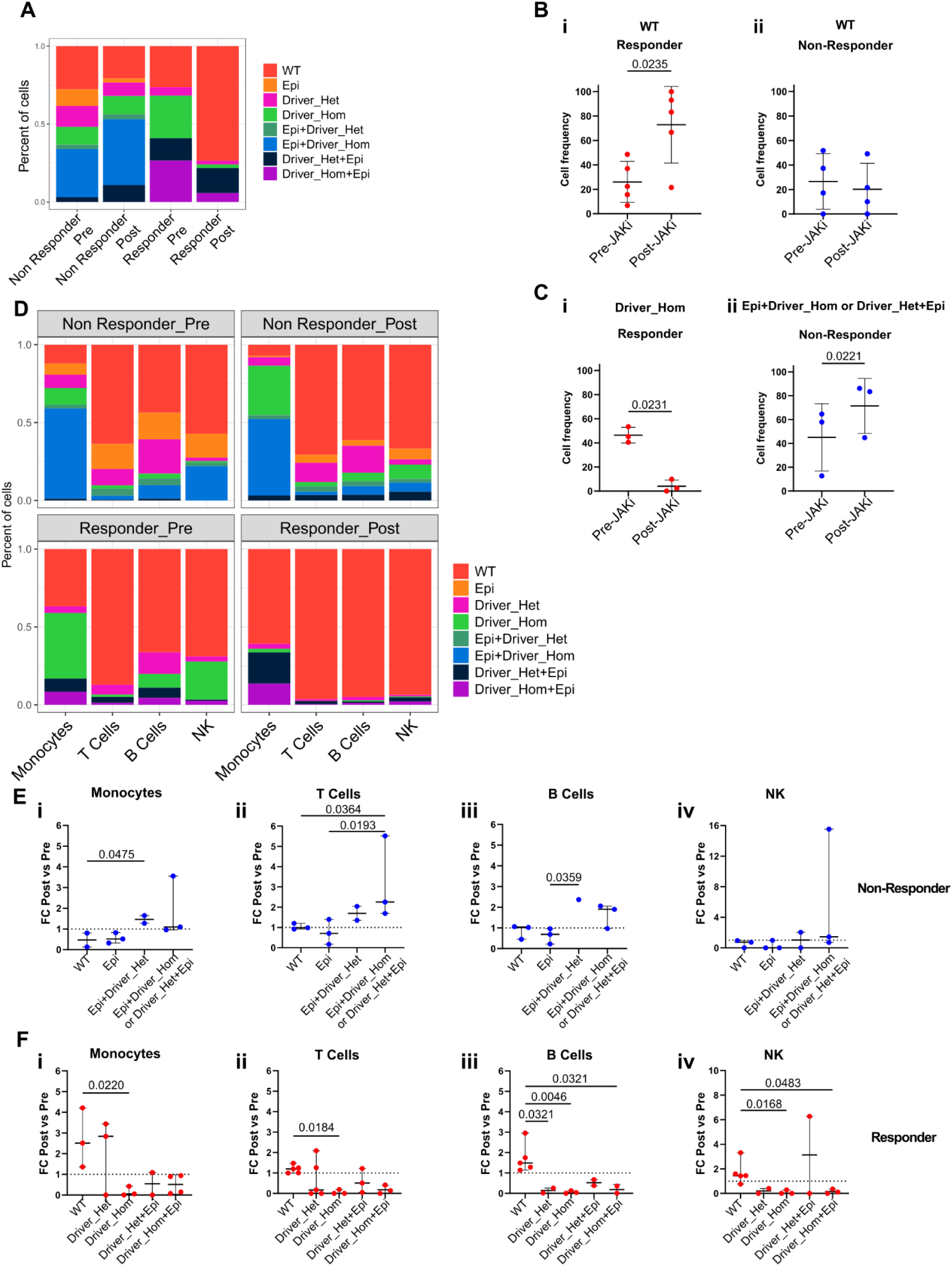
Co-mutated clones are expanded after treatment in CD34- cells from Non-Responder patients. **A** Bar plot shows the cumulative frequency of the identified genetic groups in CD34- differentiated cells classified according to timepoint and treatment response. **B** Scatter plot showing the variation of WT cells frequency in Responder (**i**) (n=5) and Non-Responder (**ii**) (n=4) patients before and after JAK inhibitor treatment. Each dot represents a patient. **C** Scatter plot showing the frequency variation of cells homozygous for MPN driver mutation in Responder patients (**i**) (n=3) or cells homozygous for MPN driver mutation and preceded by epigenetic variant (Epi+Driver_Hom) or heterozygous for MPN driver variant followed by epigenetic mutation (Driver_Het+Epi) in Non-Responder patients (**ii**) (n=3) before and after JAK inhibitor treatment. Each dot represents a patient. **D** Bar plot showing the cumulative frequency of the identified genetic groups across CD34- differentiated cell clusters. Cells were distinguished according to timepoint and treatment response. **E** In Non-Responder patients, 4 genetic groups were shared by most of the patients. Considering the different proteomic cell clusters, monocytes (**i**), T cells (**ii**), B cells (**iii**) and NK cells (**iv**), the fold change (FC) of each genetic group was calculated in each patient and shown in the scatter plot. Each dot represents a patient (n=3). **F** In Responder patients, 5 genetic groups were shared by most of the patients. Considering the different proteomic cell clusters, monocytes (**i**), T cells (**ii**), B cells (**iii**) and NK cells (**iv**), the fold change (FC) of each genetic group was calculated in each patient and shown in the scatter plot. Each dot represents a patient (n=5). Comparisons were performed using Friedman test for paired samples, and Mann-Whitney test for unpaired samples followed by uncorrected Dunn’s test. P-values less than 0.05 are considered significant and reported in panels. Abbreviations: FC: fold change. Driver_Het: clone harboring heterozygous MPN driver mutation; Driver_Hom: clone harboring homozygous MPN driver mutation; Driver_Het+Epi: clone harboring heterozygous MPN driver mutation followed by variant(s) in epigenetic regulator(s); Driver_Hom+Epi: clone harboring homozygous MPN driver mutation followed by variant(s) in epigenetic regulator(s); Epi: clone harboring mutation(s) in epigenetic regulator(s); Epi+Driver_Het: clone harboring variant(s) in epigenetic regulator(s) followed by heterozygous MPN driver mutation; Epi+Driver_Hom: clone harboring variant(s) in epigenetic regulator(s) followed by homozygous MPN driver mutation; WT: wild type clone.

### WT monocytes are expanded in patients who respond to JAK inhibition

Based on these observations we moved on by analyzing changes in clonal composition of different cellular populations. Firstly, we focused on CD34^-^ differentiated cells and performed a global analysis by splitting cells according to treatment response and timepoint. As represented by bar plots in Figure 5D clonal prevalence varied greatly within Resp cells (Fig. S4A). In particular, the T-, B-, and NK-cell compartments were the most affected as mutated clones almost disappeared after treatment. WT cells also expanded in T-, B- and NK-cells in Non-Resp; however, mutated clones persisted at a non-negligible frequency despite treatment. As regards monocytes, myeloid cells belonging to the neoplastic clone, in Resp WT clone expanded, whereas Driver_Hom clone nearly disappeared (Fig. 5D and Fig. S4B, C). Conversely, in Non-Resp, no significant changes in clone frequency were observed (Fig. 5D and Fig. S4B, C).

These variations were confirmed by analyzing clone frequency variation in different cell clusters within each patient (Fig. 5E, F). In Non-Resp WT cells frequency was almost unaffected by JAKi while co-mutated clones were significantly expanded only among monocytes and T lymphocytes (Fig. 5E). By contrast, in Resp patients a significant expansion of WT cells was observed in all cell compartments (Fig. 5F). Particularly, while in T-, B- and NK cells the frequencies of mutated clones were reduced in almost all patients, among monocytes only Driver_Hom were significantly reduced as compared to WT cells (Fig. 5F).

Collectively our results revealed that a significant clonal rewiring was observed only in Resp patients with the greatest effects restricted to lymphoid lineages. As regards the myeloid lineage WT cells were expanded at the expense of Driver_Hom ones, but co-mutated clones appeared to be expanded after JAK inhibition also in Res (Fig. 5D).

### JAKi does not eradicate co-mutated clones in the HSPC compartment

The observed effects in differentiated cell populations might represent either the result of the direct action of the JAK inhibitor on patients’ monocytes and lymphocytes, or the consequence of clonal selection occurring in stem and progenitor cell subpopulations, or the combination of both. To explore this issue, we analyzed clonal dynamics among CD34+ HSPCs.

Considering CD34+ cells as a whole it was evident that in Resp the WT clone expanded while Driver_Hom cells percentage was reduced, whereas in Non-Resp, WT cells were only a small fraction and further decreased following treatment. Notably, in both Resp and Non-Resp, clones harboring both MPN driver and epigenetic variants were nearly half of the cells prior to treatment and expanded despite JAK inhibition (Fig. 6A and Fig. S5). Per-patient analysis yielded consistent results demonstrating that co-mutated cells significantly expanded after treatment in Non-Resp, while in Resp the frequency of Driver_Hom cells was significantly reduced relative to both WT and Driver_Het+Epi cells (Fig. 6B).

**Figure 6:**
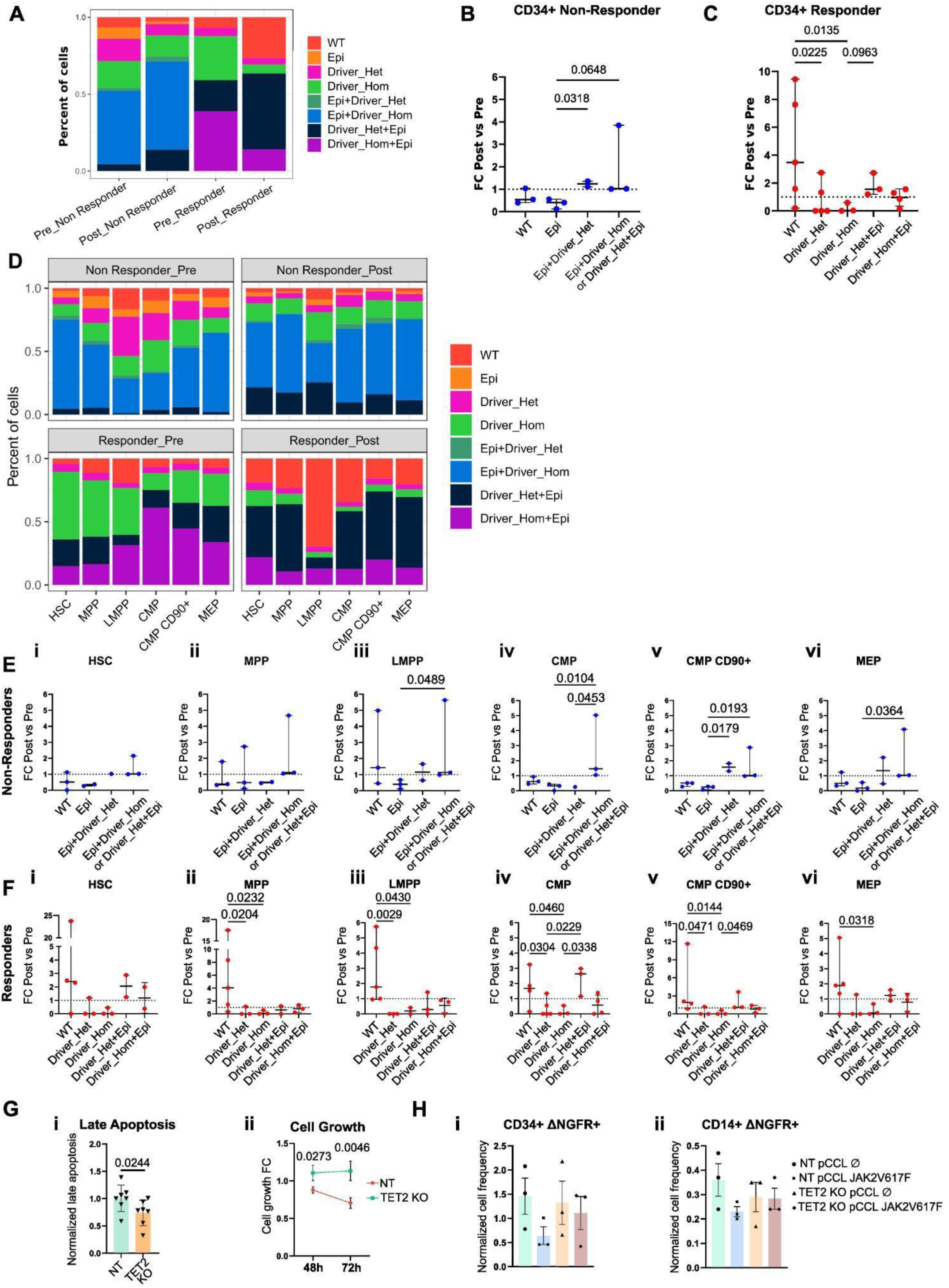
MPN driver homozygous cells frequency is reduced in CD34+ hematopoietic stem and progenitor cells (HSPCs) after treatment, while co-mutated clones persist over time. **A** Bar plot showing the cumulative frequency of the identified genetic groups in CD34+ HSPCs classified according to timepoint and treatment response. **B** In Non-Responder patients, 4 genetic groups were shared by most of the patients. The scatter plot displays the fold change (FC) of the frequency of every genetic group within CD34+ HSPCs in each Non-Responder patient. Each dot represents a patient (n=3). **C** The scatter plot displays the FC of the frequency of every genetic group within CD34+ HSPCs in each Responder patient. Each dot represents a patient (n=5). **D** Bar plot shows the cumulative frequency of the identified genetic groups across CD34+ HSPC clusters. Cells were distinguished according to timepoint and treatment response. **E** Non-Responder patient samples were analyzed by distinguishing the identified proteomic cell clusters of CD34+ HSPCs including, hematopoietic stem cells (HSC) (i), multipotent progenitors (MPPs) (ii), lymphoid-primed multipotent progenitors (LMPPs) (iii), common myeloid progenitors (CMPs) (iv), CMP positive for CD90 (CMP CD90+) (v) and megakaryocyte and erythroid progenitors (MEPs) (vi). The FC of each genetic group was calculated within each cluster in each patient and shown in the scatter plot. Each dot represents a patient (n=3). **F** Responder patient samples were analyzed by distinguishing the identified proteomic cell clusters of CD34+ HSPCs including, HSC (i), MPPs (ii), LMPPs (iii), CMPs (iv), CMP CD90+ (v) and MEPs (vi). The FC of each genetic group was calculated within each cluster in each patient and shown in the scatter plot. Each dot represents a patient (n=5). G *TET2* unmodified (NT) and *TET2* knock-out (TET2 KO) Hel cells were treated with Ruxolitinib (Ruxo) and apoptosis was evaluated using AnnexinV-Propidium iodide (PI) biparametric flow-cytometry after 48 hours of treatment. The gating strategy is shown in Supplementary Figure 6. The frequency of late apoptotic cells was reported in panel **i**. Cell growth was evaluated 48 and 72 hours after treatment (**ii**). **H** We set up an in vitro model of sequential mutation acquisition in cord blood (CB) derived CD34+ cells by introducing TET2 KO, using the same CRISPR/Cas9 strategy, and JAK2V617F lentiviral-mediated overexpression. We obtained 4 experimental conditions, control cells with unmodified TET2 and transduced with empty pCCL vector (NT pCCL ∅), cells overexpressing JAK2V617F variant with unmodified TET2 (NT pCCL JAK2V617F), TET2 KO cells transduced with empty pCCL vector (TET2 KO pCCL ∅) and TET2 KO cells overexpressing JAK2V617F variant (TET2 KO pCCL JAK2V617F). Cells were treated with Ruxo and the frequency of NGFR+ transduced cells was evaluated by means of cytofluorimetric analysis (for details see Supplementary Figure 8). Bar plots display the normalized frequency of NGFR+ CD34+ (**i**) or CD14+ (**ii**) cells in different samples (n=3). Comparisons were performed using Friedman test for paired samples, and Mann-Whitney test for unpaired samples followed by uncorrected Dunn’s test. Abbreviations: CMP: common myeloid progenitors; CMP CD90+: CMP positive for CD90; FC: fold change; HSC: hematopoietic stem cells; LMPP: lymphoid-primed multipotent progenitors; MEP: megakaryocyte and erythroid progenitors; MPP: multipotent progenitors; Ruxo: Ruxolitinib; NT: non targeting; NT pCCL∅: sample nucleofected with CRISPR-Cas9 Non Targeting guide and transduced with empty pCCL lentiviral vector; NT pCCL JAK2V617F: sample nucleofected with CRISPR-Cas9 Non Targeting guide and transduced with pCCL JAK2V617F lentiviral vector; TET2 KO pCCL∅: sample nucleofected with CRISPR-Cas9 *TET2*-targeting guide and transduced with empty pCCL lentiviral vector; TET2 KO pCCL JAK2V617F: sample nucleofected with CRISPR-Cas9 *TET2*-targeting guide and transduced with pCCL JAK2V617F lentiviral vector; ΔNGFR+: cells expressing Nerve Growth Factor Receptor; Driver_Het: clone harboring heterozygous MPN driver mutation; Driver_Hom: clone harboring homozygous MPN driver mutation; Driver_Het+Epi: clone harboring heterozygous MPN driver mutation followed by variant(s) in epigenetic regulator(s); Driver_Hom+Epi: clone harboring homozygous MPN driver mutation followed by variant(s) in epigenetic regulator(s); Epi: clone harboring mutation(s) in epigenetic regulator(s); Epi+Driver_Het: clone harboring variant(s) in epigenetic regulator(s) followed by heterozygous MPN driver mutation; Epi+Driver_Hom: clone harboring variant(s) in epigenetic regulator(s) followed by homozygous MPN driver mutation; WT: wild type clone.

In different subpopulations of HSPCs JAK inhibition failed to counteract clonal expansion of most mutated clones in Non-Resp cells; indeed, Epi+Driver_Het and Driver_Het+Epi clones became predominant across the different HSPC subpopulations. Notably, HSC clonal composition underwent only minor changes as it was already dominated by co-mutated clones prior to JAK inhibition. In Resp, the WT clone expanded in every cell cluster at the expense of Driver_Hom cells, nevertheless, Driver_Het+Epi clone expanded as well. Only in LMPP Resp cells the WT clone dramatically expanded after JAK inhibition and became prevalent (Fig. 6D). Accordingly, per-patient analysis demonstrated that WT cells significantly expanded compared with Driver_Hom cells in Resp across all cell clusters, except for HSCs. Nevertheless, co-mutated clones also significantly expanded in both Resp and Non-Resp within myeloid progenitor compartments (CMP, CMP CD90+ and MEP), suggesting the presence of a specific therapy-resistance trait in these clones (Fig. 6E, F).

### JAK2V17F single mutated cells are more sensitive to JAK inhibition

According to our SC multiomic findings, JAK2V617F homozygous clones appear to be the most sensitive to JAK inhibition. On the other hand, co-mutated clones, both in Resp and Non-Resp appeared to be more resistant to JAKi as their frequency increased over time. To validate our results *in vitro* we developed genetically engineered cell models by introducing *TET2* knock-out (KO) in JAK2V617F homozygous Hel cells and cord blood (CB) derived CD34^+^ cells using CRISPR/Cas9. Preliminary experiments allowed us to select the most effective guide RNA (gRNA) to obtain *TET2* KO and abrogation of its demethylase activity (Fig. S6A-D). Co-mutated *TET2* KO Hel cells were more resistant to JAK inhibition compared to unmodified JAK2V617F homozygous cells as demonstrated by reduced apoptosis induction and increased cell numbers (Fig. 6G and Fig. S6E). Accordingly, edited TET2 allele did not vary upon Ruxo treatment suggesting the resistance of co-mutated cells to JAK inhibition (Fig. S6F).

Given that Non-Resp displayed more frequently the acquisition of MPN driver mutation as a secondary event, we set up an *in vitro* model in CB derived CD34+ cells by sequentially introducing *TET2* KO and JAK2V617F lentiviral-mediated overexpression (Fig. S7). Genetically engineered CD34+ cells were subjected to Ruxo treatment at two different timepoints, right after immunomagnetic cell selection, when most cells belonged to the hematopoietic stem cell compartment, and after 3 days of *in vitro* culture, when myeloid differentiation had already been induced. After treatment the frequency of ΔNGFR+ transduced cells was evaluated among CD34+ HSPC and CD14+ monocyte cell fractions respectively (Fig. S8A). I*n vitro* results confirmed that NT pCCL JAK2V617F cells are the most sensitive to pharmacologic JAK inhibition and *TET2* KO provides therapy resistance also in the presence of the MPN driver mutation (Fig. 6H and Fig. S8B).

## Discussion

MF arises from the acquisition of driver somatic mutations in HSCs that lead to the constitutive activation of the JAK/STAT signaling pathway. These mutations affect *JAK2*, *MPL* and *CALR* genes and are detected in almost 90% of patients^28^. Soon after the identification of JAK2V617F, the most common driver event, the first-in-class JAK1/2 inhibitor Ruxolitinib was identified and approved for the treatment of MF^12^. Since then, several molecules have been tested leading to more recent approvals of other JAKi with different specificity including Fed, Pacritinib and Momelotinib^29^. While JAKi are effective in reducing spleen volume and improving quality of life, their impact on tumor burden is limited; consequently, they do not prevent disease progression or leukemic transformation. According to clinical trials, Ruxo and Pacritinib achieved a median reduction in JAK2V617F allele burden of 8% and 15.8%, respectively^30,31^, whereas Fed failed to induce meaningful changes^32^. Nevertheless, a greater molecular response has been reported in patients with a higher baseline JAK2V617F allele burden treated with Ruxo and Fed^33–35^.

To improve our understanding of the molecular response to JAKi in MF, we analyzed the clonal architecture and dynamics at the SC level in a cohort of 12 JAKi treated patients, stratified according to their clinical outcome. Bulk analysis failed to reveal significant variation in MPN driver mutation VAF following JAK inhibition, while SC genomics provided meaningful insights into the extent of molecular response and its determinants. Indeed, in Resp MPN driver mutation was the first or the sole acquired somatic event, whereas in Non-Resp it was more frequently preceded by the acquisition of a non-driver mutation affecting epigenetic modifier genes, particularly *TET2* and *ASXL1*. This is in line with previous findings demonstrating that Ruxo impairs the clonogenicity of mutated CD34^+^ cells from JAK2-first patients, while cells from TET2-first patients are not affected^2^. These observations indicate that patients who had acquired an MPN driver mutation after a variant in epigenetic modifier genes are less sensitive to JAKi and at higher risk of leukemic transformation. Indeed, leukemic transformation can occur through the expansion of specific subclones already present during the chronic phase of the disease in patients displaying the driver mutation as a secondary event, as is the case of patient #4. These clones can be either subclonal or phylogenetically independent to the one dominating the chronic phase^10,11^.

Compared to bulk analysis and colony based approaches, SC techniques allow a better reconstruction of clonal architecture and dynamics^36^ and the simultaneous inquiry into multiple information layers^11,37,38^. By performing coupled SC genomic and proteomic analysis, we demonstrated that JAKi response was associated with the expansion of WT cells and contraction of most mutated clones only in Resp in both CD34^+^ and CD34^-^ compartments. As expected, the expansion of WT cells in Resp was much more evident among differentiated cells, which include lymphoid cells that are commonly known to be devoid of MPN driver mutations^39^. Our results clearly suggest the presence, in both Resp and Non-Resp, of a minor proportion of mutated cells within lymphoid clusters, according to recent observations demonstrating that clonal hematopoiesis can contribute to B cell and to a lesser extent T cell lymphopoiesis^40,41^. The effect of JAKi treatment distinguishes Resp and Non-Resp because only in Resp, mutated clones almost completely disappeared from lymphoid clusters and are significantly reduced among LMPPs. As regards monocytes, Non-Resp and Resp profoundly differ as only in Resp WT cells expanded even if mutated clones did not disappear. This finding is consistent with the limited modulations in MPN driver VAF observed in granulocytes using bulk NGS.

SC multiomic analysis provided important clues about the causes of the higher sensitivity to JAKi in patients harboring the MPN driver variant as the founder mutation. Indeed, only these patients display clones harboring the sole homozygous MPN driver mutation, particularly JAK2V617F. These cells were the most affected by JAK inhibition, and their frequency was significantly reduced after treatment. Conversely, in the same patients, heterozygous and co-mutated clones were not affected by JAKi. Indeed, co-mutated clones take advantage of the elimination of homozygous JAK2V617F clones and expand after treatment. This effect is exemplified by the clonal dynamics observed in patient #8 harboring the duplication of the 9p chromosome which encodes for *JAK2*^27^. JAK inhibition leads to the expansion of the clone displaying 9p duplication and only two *JAK2* mutated alleles in both CD34+ and CD34- cells. The elimination of the sole homozygous MPN driver clone might represent a justification for the limited capacity of identifying a molecular response in patients, based on conventional bulk genomic analysis. At the same time, this emphasizes the effectiveness of JAK inhibition and suggests that only homozygous JAK2V617F clones are dependent on JAK/STAT oncogenic signaling. This interpretation is consistent with data coming from clinical practice because Ruxo is much more effective in terms of molecular response in PV and PPV-MF patients^42–44^ characterized by higher frequency of JAK2V617F homozygosity and, as regards PV, reduced incidence of secondary non-driver mutations^1,45^. As proposed by Ross and colleagues^46^ we might speculate that our observation is the result of two concurring effects. On one hand JAK2V617F homozygous cells appear to be more dependent on oncogenic signaling in line with recent data demonstrating that CRISPR/Cas9 based JAK2V617F inactivation eradicates only homozygous cells while heterozygous ones maintain viability and differentiation potential^47^. On the other hand, the presence of additional somatic mutations reduces cell dependency on JAK2V617F oncogenic signal as demonstrated by our *in vitro* models, which clearly show that the combination of mutations, particularly JAK2V617F and *TET2 KO*, reduces the sensitivity of cells to Ruxo. The JAKi refractoriness of co-mutated clones in both Non-Resp and Resp results in their persistence through treatment, particularly within HSPCs clusters, where WT cells always represent a smaller fraction of cells. Given that, it is noteworthy that HSPCs remain as a disease reservoir after treatment not only in Non-Resp but also in Resp when driver mutation is accompanied by additional somatic variants.

Collectively, our results suggest that the order of mutation acquisition is one of the primary determinants of JAKi response in MF patients, because JAKi suppresses the expansion of homozygous MPN driver clones but has only limited effects on co-mutated ones. Indeed, only patients who have acquired the MPN driver mutation as the first somatic event can present with a homozygous MPN driver clone that is targeted by JAKi and whose contraction might be reflected by the VAF decrease. Given that bulk genomic analysis cannot resolve the presence and relative abundance of homozygous and heterozygous clones, the power of mutation acquisition order inference is limited. Therefore, SC genomic analysis represents a more powerful and reliable tool for the characterization of clonal architecture in MPN patients that might allow the identification of patients who would benefit the most from JAK inhibition. On the other hand, our results demonstrate that co-mutated clones may benefit from JAKi treatment by outcompeting other neoplastic cell populations and persist as a disease reservoir within the stem and progenitor cell compartments in both Resp and Non-Resp. Further investigations are needed to identify novel therapeutic strategies targeting co-mutated clones that can boost molecular response in MF overcoming clones therapy resistance.

## Methods

### Human samples

The study included 12 patients diagnosed with primary or secondary MF, under treatment in the Hematology department of Azienda Ospedaliera-Universitaria of Modena. Primary MF was diagnosed according to 2016 WHO criteria^48^, while secondary MF was defined based on the IWG-MRT criteria^49^. PBMCs were isolated via density-gradient centrifugation using Lympholyte-H (Cedarlane, Burlington, Canada), while CD34⁺ cells were immunomagnetically sorted using CD34 MicroBead Kit UltraPure (Miltenyi Biotec, Bergisch Gladbach, Germany). Granulocytes were purified by red blood cell lysis using an ammonium chloride-based lysis buffer. Human CD34⁺ cells were purified from umbilical CB samples, collected after normal deliveries, according to the institutional guidelines for discarded material.

The study was conducted in compliance with the Declaration of Helsinki under the local Institutional Review Board’s approved protocol. All subjects involved in the study provided informed written consent.

### Bulk NGS analysis

As previously described^11^, gene mutations were detected in genomic DNA (gDNA) extracted from whole peripheral blood using a capture-based target enrichment approach with the CE-IVD Myeloid Solution™ kit (SOPHiA Genetics, Rolle, Switzerland). Libraries were sequenced on an Illumina MiSeq platform (Illumina, San Diego, CA, USA).

### Single-cell DNA+Protein analysis

PBMCs and HSPCs from MPN patients were thawed following 10× Genomics® “Sample Preparation Demonstrated Protocol” (10× Genomics, Pleasanton, CA, USA) and mixed in a 1:1 ratio.

Cells were labelled with an Antibody-Oligo Conjugated (AOCs) cocktail (TotalSeq™-D Human Heme Oncology Cocktail, BioLegend, San Diego, California, USA) supplemented with additional antibodies (**Table S3**). Next, cells underwent single-cell DNA+Protein library preparation following manufacturer’s instructions. In detail, barcoded samples were subjected to targeted PCR amplification through a custom 299-amplicon DNA panel (**Table S4**). Pooled purified libraries were sequenced using a 2x150bp paired-end chemistry on a NovaSeqX platform (service by Macrogen s.r.l, Milan, Italy). Resulting FASTQ files were pre-processed on Tapestri Pipeline v2.

Variant selection was performed using Tapestri Insights v.2.2 software, prioritising variants previously identified in bulk sequencing and selecting those classified as pathogenic/likely pathogenic according to the ACMG criteria^50^. Individual .h5 files were processed through Mosaic v3.12.2 to assess CNVs and generate raw protein expression matrices. Cells harboring both genomic and proteomic information were retained for downstream analyses. Cohort-level analyses were conducted using Seurat R package (RRID:SCR_016341). The Seurat object was generated by loading the raw protein expression matrices in the *counts* slot, while genotype was added as *metadata*. Raw protein counts were Log-normalized, and cells with IgG signal <0,1 were retained. Protein expression was then scaled, followed by Principal Component Analysis (PCA), nearest neighbor construction and UMAP.

Cell clustering based on protein expression was performed using *FlowSOM* package (RRID:SCR_016899) through an iterative approach. Raw protein counts were normalized using Centered Log-Ratio (CLR) transformation and initially subset to include markers associated with lineage-positive cells and CD34 antigen. Through this approach we identified multiple differentiated cell populations and a cluster comprising CD34⁺ cells, which was subsequently sub-clustered using HSPCs-associated markers. Cluster annotation was performed based on published literature data^10^.

To avoid over-representation of individual samples within the cohort, annotated cells were downsampled to 1000 cells/sample while preserving the original intra-sample distribution of annotated cell clusters. The similarity between genotype distributions in the full dataset and the downsampled subset was verified prior to downstream analyses.

Protein expression data and associated metadata were visualized using *Seurat*, *ggplot2* (RRID:SCR_014601), *dittoseq*^51^ and *MiloR* (RRID:SCR_025630) packages. Mutational data were plotted with *ClevRvis* (RRID:SCR_023154) and *timescape* R packages^52^. Hierarchical clustering of protein markers was performed through hclust function on scaled average expression data.

### CRISPR/Cas9-based Genome Engineering

TET2 crRNAs and tracrRNA (sgRNA sequences are listed in **Table S5**) were resuspended in nuclease-free duplex buffer (IDT Integrated DNA Technologies, Coralville, IA, USA) at a concentration of 200μM and annealed by mixing equimolar amounts to obtain a final concentration of 100pmol/μl. Ribonucleoprotein complexes (RNPs) were generated by incubating 120pmol of annealed crRNA:tracrRNA with 105pmol of Alt-R HiFi Cas9 Nuclease V3 (IDT) for 10’ at room temperature (RT). Subsequently, 1μl Alt-R Cas9 Electroporation Enhancer (IDT) was added. Next, 1x10^6^ CB CD34⁺ or Hel cells were resuspended in 100μL P3 or SF solution (Lonza Ltd., Basel, Switzerland) respectively, mixed with 6μL of Alt-R RNP complexes, and electroporated using Lonza 4D-Nucleofector system (program DZ-100). gDNA was extracted as previously described^27^. The genomic regions flanking gRNA target sites were amplified by PCR using Platinum SuperFi DNA Polymerase (Thermo Fisher Scientific, Waltham, MA, USA) and the primers listed in **Table S5**.

Editing efficiency was assessed by tracking of indels by decomposition (TIDE) software^58^ on PCR amplicons of the genomic region surrounding the gRNA target site in treated cells (Sanger sequencing service by Microsynth AG, Balgach, Switzerland).

### JAK2V617F overexpression

A full-length human JAK2V617F cDNA was generated by reverse transcription-PCR (RT-PCR) using M-MLV reverse transcriptase (Thermo Fisher) on total RNA extracted from Hel cell line (primers listed in **Table S6**, all from IDT).

To generate pCCL-mCMV-JAK2V617F-ΔNGFR (pCCL-JAK2V617F), the JAK2V617F cDNA cassette was inserted into pCCL-CMV-GFP-ΔNGFR (pCCL-EV) lentiviral vector to replace the GFP cassette^53^. Briefly, fragments for the minimal CMV promoter, JAK2V617F (1-1729 nt) and JAK2V617F (1689-3399 nt) were amplified by PCR with Platinum SuperFi II PCR Master Mix (ThermoFisher) and primers in **Table S6**. PCR amplicons were inserted by Gibson Assembly in pCCL-EV lentiviral vector, previously digested with *EcoRV* and *NheI* (New England Biolabs, Ipswich, MA, USA) to remove the GFP cassette. The resulting plasmid was confirmed by Whole Plasmid Sequencing (service by Eurofins Scientific, Luxembourg, Luxembourg).

Lentiviral particles carrying either pCCL-EV or pCCL-JAK2V617F were produced in HEK-293T cells as previously described^53^.

Lentiviral transduction of CB CD34⁺ cells was performed in “Stem” culture medium (**Table S7**) as previously reported at MOI=1^54^.

Two days after the infection, transduced cells were immunomagnetically purified using the EasySep™ “Do-It-Yourself” Selection Kit (STEMCELL Technologies, Vancouver, BC, Canada) using the mouse anti-human p75-NGFR (BD Biosciences, Franklin Lakes, NJ, USA) as previously described^10^.

### Cell culture

CB CD34⁺ cells were engineered as follows. Upon thawing, cells were seeded 5x10^5^ cells/ml “Stem” culture medium (**Table S7**) for 2 days. Cells were then nucleofected either with TET2-targeting gRNA or Non Targeting (NT) gRNA. After four days, cells were transduced either with pCCL-EV or pCCL-JAK2V617F. Following immunomagnetic selection, ΔNGFR⁺CD34⁺ cells were seeded 5x10^5^ cells/ml in “Multilineage” culture medium (**Table S7**).

HEK-293T cells were cultured as previously described^53^.

Hel cell line was seeded at 3x10^5^ cells/ml and maintained in RPMI-1640 supplemented with 1X Glutamine, 1X Penicillin/Streptomycin (all from Euroclone) and 10% FBS (ThermoFisher Scientific).

### Ruxolitinib treatment

Engineered CD34⁺ cells and Hel cells were treated daily for 4 days with Ruxolitinib 2,5μM or 1μM respectively. Apoptosis was assessed after 48 hours by AnnexinV-PI staining and cell growth was evaluated after 48 hours and 72 hours of treatment.

### Flow Cytometry

Following Ruxolitinib treatment, engineered CD34⁺ cells were incubated with “FcR Blocking Reagent, human” (Miltenyi Biotec) and stained with antibodies against CD34, CD14, CD15 and CD271 (LNGFR) for 20’ at 4°C in the dark. Full details on antibodies are reported in **Table S8**.

AnnexinV-PI staining on Hel cells was performed as previously described^55^.

Flow cytometric analyses were performed using a BD FACSCanto II or BD FACSymphony A1 (BD Biosciences). Data were analyzed using FlowJo_v10.10.0 software. Representative gating strategies for CD34⁺ cells and HEL cells are shown in **Figures S8A** and **S6E**, respectively.

### Dot Blot

300ng of gDNA were denatured 5’ at 95°C, rapidly cooled in ice, and spotted onto a Amersham™Hybond-N+ nylon membrane (Cytiva, Marlborough, MA, USA). After air-drying, DNA was UV cross-linked for 3’ and subsequently blocked in 0,1% TBS-T + 5% non-fat milk. The membrane was hybridized overnight with the primary and secondary antibodies listed in **Table S8**. Chemiluminescence signals were detected using ECL Select Western Blotting detection reagent (Sigma-Aldrich) through ChemiDoc™ MP Imaging System (Bio-Rad Laboratories Inc., Hercules, CA, USA). To normalize signal intensities, DNA loading was assessed by staining the membrane for 10’ with 0,2% methylene blue in 0,3% sodium acetate.

### Western Blot

Total protein extracts were obtained by RIPA buffer and loaded onto 7,5% SDS-Polyacrilamide gels as previously described. Membranes were immunoblotted with the primary and secondary antibodies listed in **Table S8**. Chemiluminescent signals were detected using ECL Select Western Blotting detection reagent (Sigma-Aldrich) through ChemiDoc™ MP Imaging System.

### ddPCR

To assess JAK2V617F overexpression in transduced CD34+ cells, total RNA was extracted using the miRNeasy Micro Kit (QIAGEN) and 100 ng were retrotranscribed using High-Capacity cDNA Reverse Transcription Kit (ThermoFisher Scientific), as previously described^27^. The resulting cDNA was analysed by droplet digital PCR (ddPCR) by using sequence-specific primers and probes, whose sequences are reported in **Table S9** (IDT). Droplets were generated using the QX200™ Droplet Generator, and PCR was carried out on the QX200™ Droplet Digital PCR System (Bio-Rad Laboratories Inc.), with an annealing temperature of 61°C. Fluorescence was measured using the QX200^TM^ Droplet Reader, and positive droplets were identified using the manufacturer’s analysis software (Bio-Rad Laboratories Inc.).

### Statistics and Data Availability

Sequencing data will be deposited in Gene Expression Omnibus (GEO). Accession numbers will be made available upon completion of the deposition process.

All bioinformatic analyses were conducted using R (version 4.3.2) and R Studio (version 2025.09.2) software. Statistical tests were performed in RStudio or Prism **10** (GraphPad Software) and are detailed in each figure panel.

## Acknowledgment

This work was supported by the following grants from: Associazione Italiana per la Ricerca sul Cancro (AIRC): AIRC 5 per 1000 project #21267, IG project #29077, MFAG project #32672; the Italian Ministry of University and Research (PRIN 2022 project #2022F4WMR3 and PRINPNRR 2022 project #P202259EM5). The research leading to these results has received funding from the European Union—NextGenerationEU through the Italian Ministry of University and Research under PNRR—M4C2-I1.3 Project PE_00000019 ‘HEAL ITALIA’ to Rossella Manfredini and Sebastiano Rontauroli (CUP E93C22001860006, University of Modena and Reggio Emilia, Modena, Italy). The views and opinions expressed are those of the authors only and do not necessarily reflect those of the European Union or the European Commission. Neither the European Union nor the European Commission can be held responsible for them.

