## Supplementary Figures for "Therapy-Induced Clonal Selection as a Driver of Response to JAK Inhibitors in Myelofibrosis"

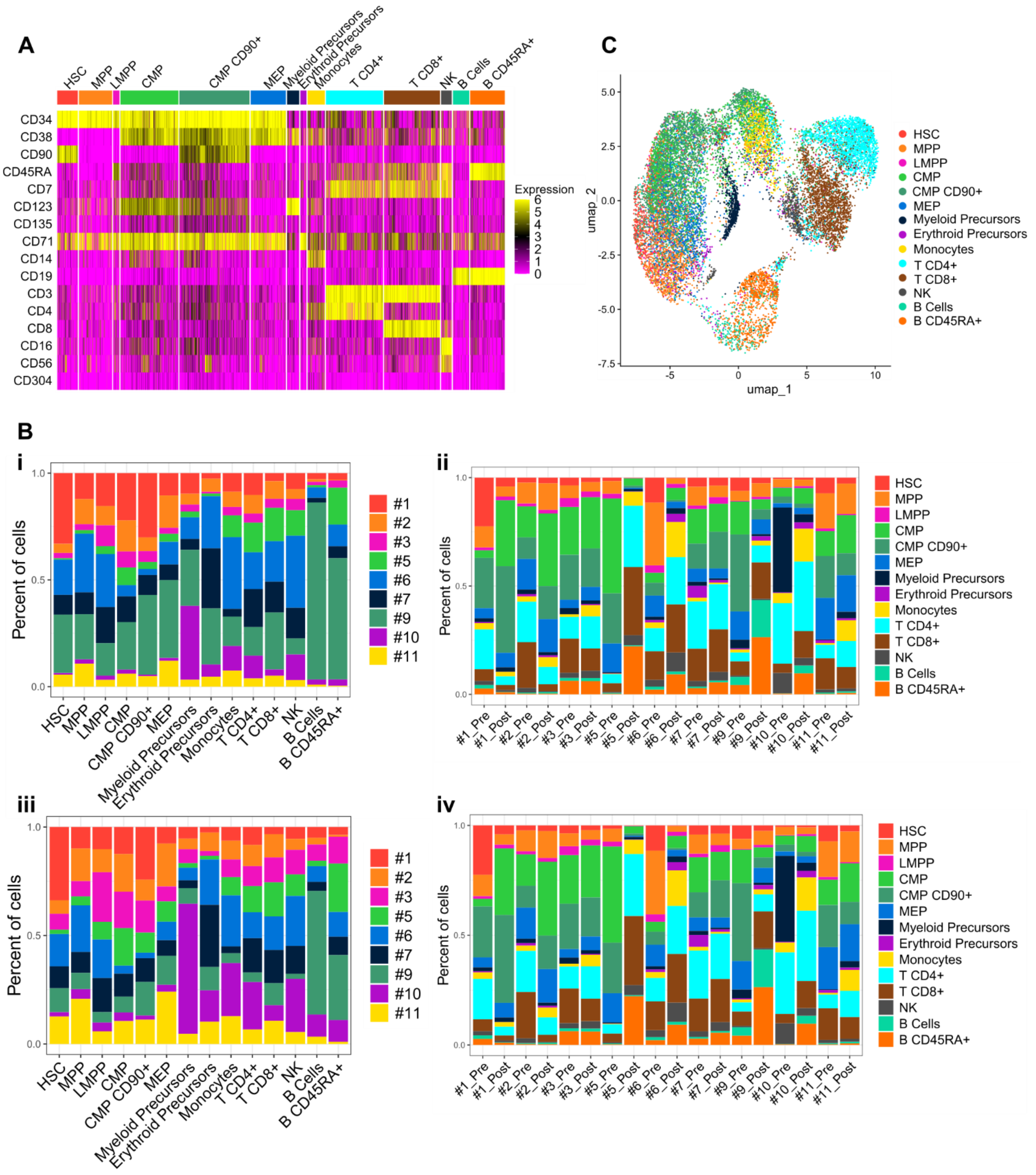


**Supplementary Figure 1: A** The heatmap shows the expression of selected antigens in cell clusters identified using FlowSOM R package. **B** Due to the variable number of recovered cells in each analyzed sample (Supplementary Table 2), in order to mitigate analysis biases due to the over-representation of specific specimens, we performed a sub-sampling of cells by including 1,000 cells per sample maintaining the relative frequency of each cell cluster within each specimen observed in the full dataset. Bar plots display the frequency of cells from different patients before (i) and after (iii) subset. Bar plots ii and iv show the frequency of hematopoietic cell clusters within each sample before (ii) and after (iv) subset. **C** UMAP representation of analysed cells after subset (n= 18000) distributed according to surface antigen expression and coloured according to cell clusters identified using FlowSOM. Abbreviations: B CD45RA+: B cells expressing CD45RA marker; CMP: common myeloid progenitors; HSC: hematopoietic stem cells; LMPP: Lymphoid primed multipotent progenitors; MEP: megakaryocyte erythroid progenitors; MPP: multipotent progenitors; NK: natural killers.

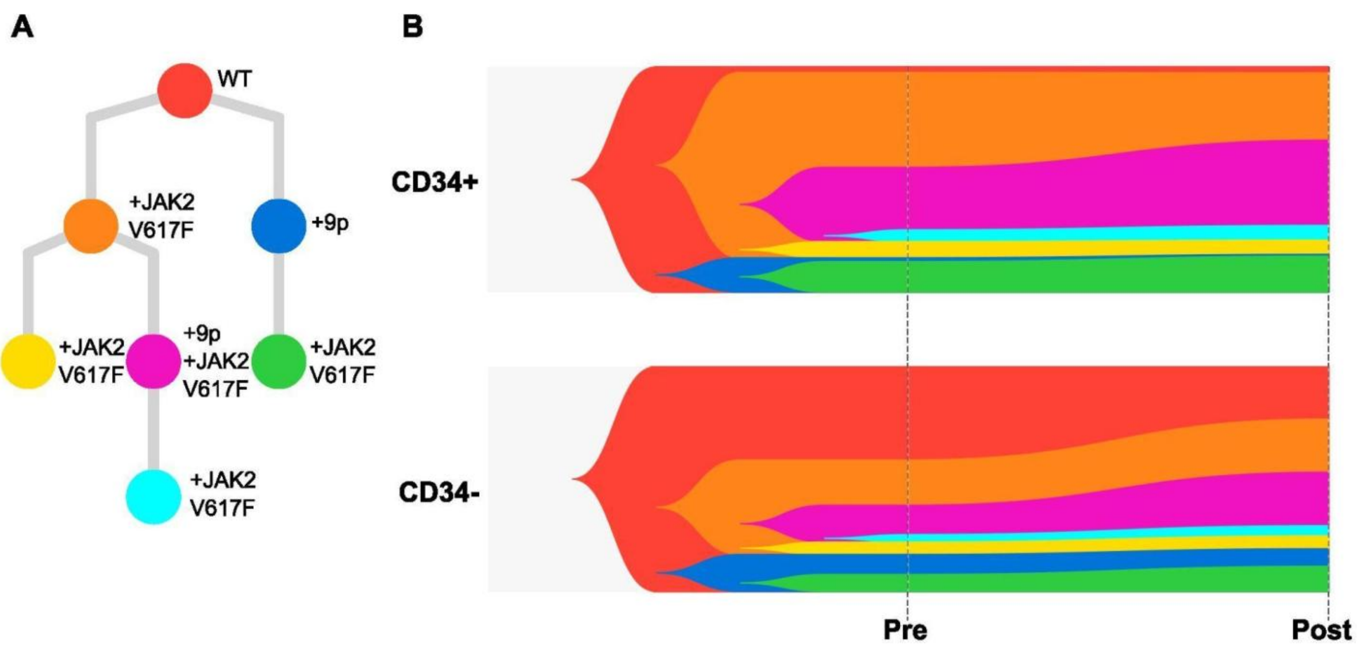


**Supplementary Figure 2**: **A** Phylogenetic tree represents the mutation and copy number variation acquisition order in patient #8 reconstructed according to single cell (SC) genomic information. The mutational event leading to the appearance of a new clone is indicated by the symbol of the affected gene. **B** Fish plot representing the clonal dynamics over time in patient #8. CD34+ hematopoietic stem and progenitor cells (HSPCs) and CD34- differentiated cells were distinguished based on SC proteomic information. Clone colors are the same used in phylogenetic tree. Abbreviations: WT: wild type; +9p: acquisition of chromosome 9 short harm duplication; +JAK2V617F: acquisition of JAK2V617F variant.

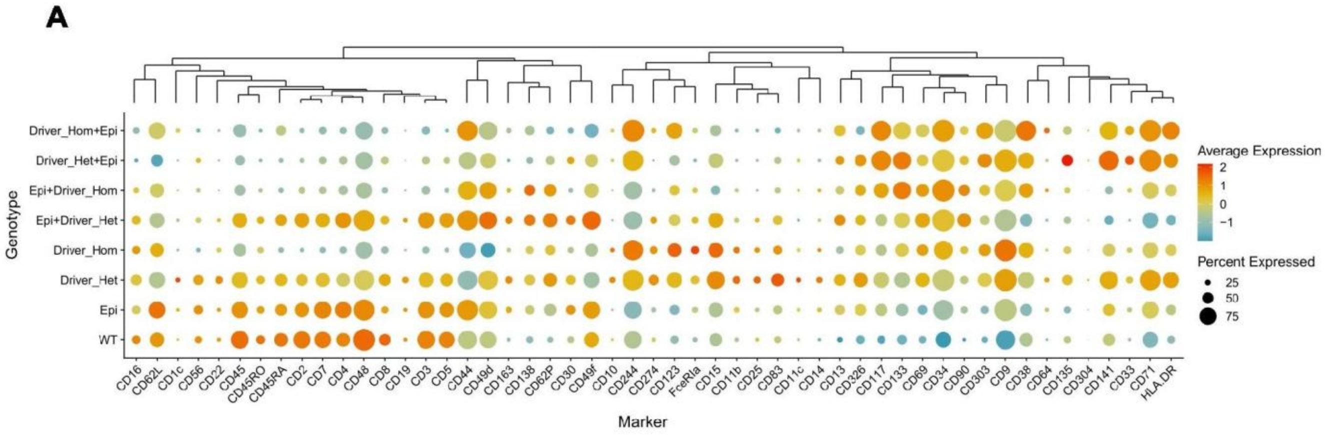


**Supplementary Figure 3**: The bubble chart displays average protein expression and frequency of cells expressing surface markers. Cells were classified according to genomic profile. Abbreviations: Driver_Het: clone harboring heterozygous MPN driver mutation; Driver_Hom: clone harboring homozygous MPN driver mutation; Driver_Het+Epi: clone harboring heterozygous MPN driver mutation followed by variant(s) in epigenetic regulator(s); Driver_Hom+Epi: clone harboring homozygous MPN driver mutation followed by variant(s) in epigenetic regulator(s); Epi: clone harboring mutation(s) in epigenetic regulator(s); Epi+Driver_Het: clone harboring variant(s) in epigenetic regulator(s) followed by heterozygous MPN driver mutation; Epi+Driver_Hom: clone harboring variant(s) in epigenetic regulator(s) followed by homozygous MPN driver mutation; WT: wild type clone.

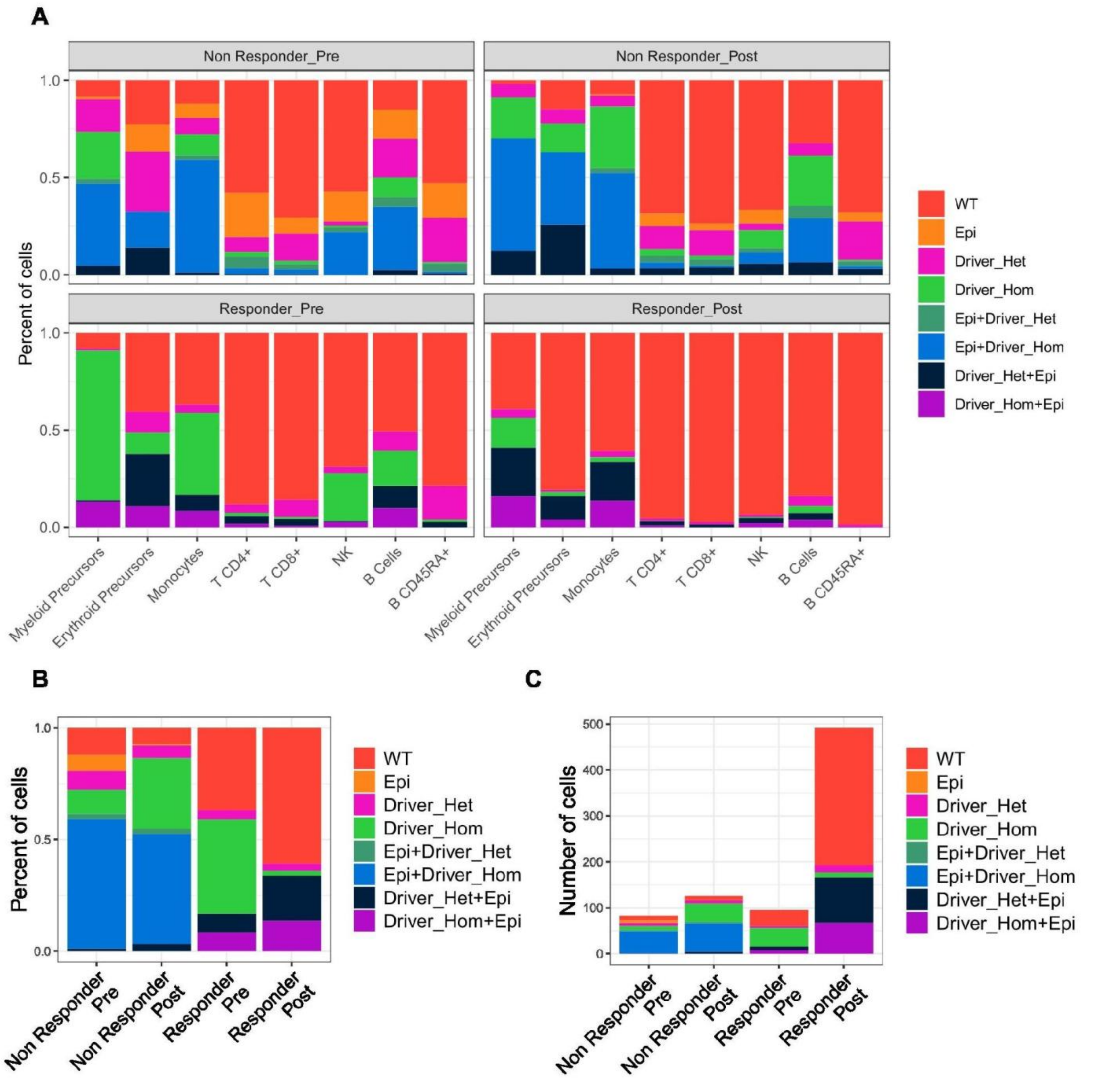


**Supplementary Figure 4**: **A** Cells were distinguished according to timepoint and treatment response in four groups. The bar plot displays the frequency of cell types within each group. **B** Bar plot showing the cumulative frequency of the identified genetic groups in monocytes. Cells were distinguished according to timepoint and treatment response. **C** Bar plot showing the absolute number of monocytes before and after JAK inhibitor treatment in Responder and non-Responder patients classified according to genetic group. Cells were classified according to timepoint and treatment response. Abbreviations: Driver_Het: clone harboring heterozygous MPN driver mutation; Driver_Hom: clone harboring homozygous MPN driver mutation; Driver_Het+Epi: clone harboring heterozygous MPN driver mutation followed by variant(s) in epigenetic regulator(s); Driver_Hom+Epi: clone harboring homozygous MPN driver mutation followed by variant(s) in epigenetic regulator(s); Epi: clone harboring mutation(s) in epigenetic regulator(s); Epi+Driver_Het: clone harboring variant(s) in epigenetic regulator(s) followed by heterozygous MPN driver mutation; Epi+Driver_Hom: clone harboring variant(s) in epigenetic regulator(s) followed by homozygous MPN driver mutation; WT: wild type clone; B CD45RA+: B cells expressing CD45RA marker; NK: natural killers.

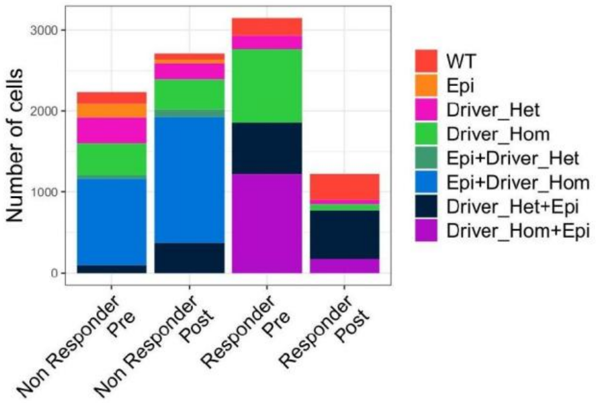


**Supplementary Figure 5**: Bar plot showing the absolute number of CD34+ hematopoietic stem and progenitor cells (HSPCs) before and after JAK inhibitor treatment in Responder and Non-Responder patients classified according to genetic group. Cells were grouped according to timepoint and treatment response. Abbreviations: Driver_Het: clone harboring heterozygous MPN driver mutation; Driver_Hom: clone harboring homozygous MPN driver mutation; Driver_Het+Epi: clone harboring heterozygous MPN driver mutation followed by variant(s) in epigenetic regulator(s); Driver_Hom+Epi: clone harboring homozygous MPN driver mutation followed by variant(s) in epigenetic regulator(s); Epi: clone harboring mutation(s) in epigenetic regulator(s); Epi+Driver_Het: clone harboring variant(s) in epigenetic regulator(s) followed by heterozygous MPN driver mutation; Epi+Driver_Hom: clone harboring variant(s) in epigenetic regulator(s) followed by homozygous MPN driver mutation; WT: wild type clone

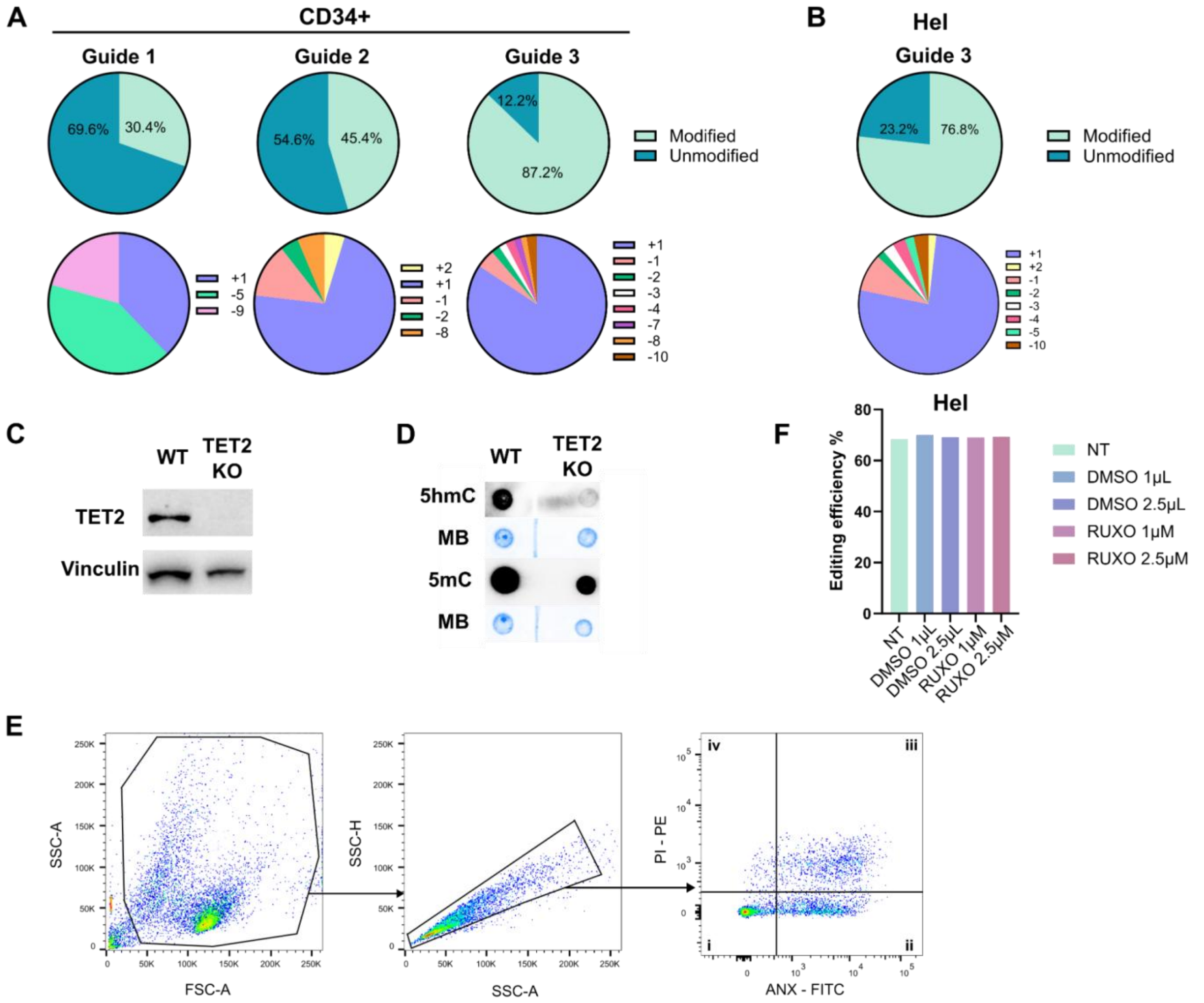


**Supplementary Figure 6**: **A** Comparison of *TET2* targeting efficiency by three different gRNAs in CD34+ cells. For each guide, the frequency of resulting modified and unmodified alleles is shown, as well as the frequency of significant in-dels detected in CD34+ cells. **B** *TET2* targeting efficiency in Hel cells using Guide 3. Pie charts show the frequency of modified and unmodified alleles and the prevalence of different in-dels generated. **C** Western blot evaluating TET2 protein expression in Hel cell line. **D** Dot blot analysis of genomic 5hmC and 5mC in unmodified and TET2 KO Hel cells. **E** Bar plot showing the editing efficiency in Hel cells after Ruxolitinib (RUXO) treatment. **F** Gating strategy used for the Annexin-V/PI apoptosis analysis. Hel cells were gated on forward (FSC) versus side scatter (SSC). Next, cells were gated on SSC-Area (SSC-A) vs SSC-Heigh (SSC-H) to exclude doublets. Then, HEL were analyzed for the ANX versus PI (A-D). Description of Panel A: live cells; B: early-stage apoptotic cells; C: apoptotic cells; D: necrotic cells or mechanically damaged cells. Abbreviations: 5hmC: 5-Hydroxymethylcytosine; 5mC: 5-Methylcytosine; ANX: Annexin V; DMSO: Dimethyl Sulfoxide; FSC-A: forward scatter-Area; MB: Methylene Blue; PI: Propidium Iodide; RUXO: Ruxolitinb; SSC-A: side scatter-Area; SSC-H: side scatter-Heigh.

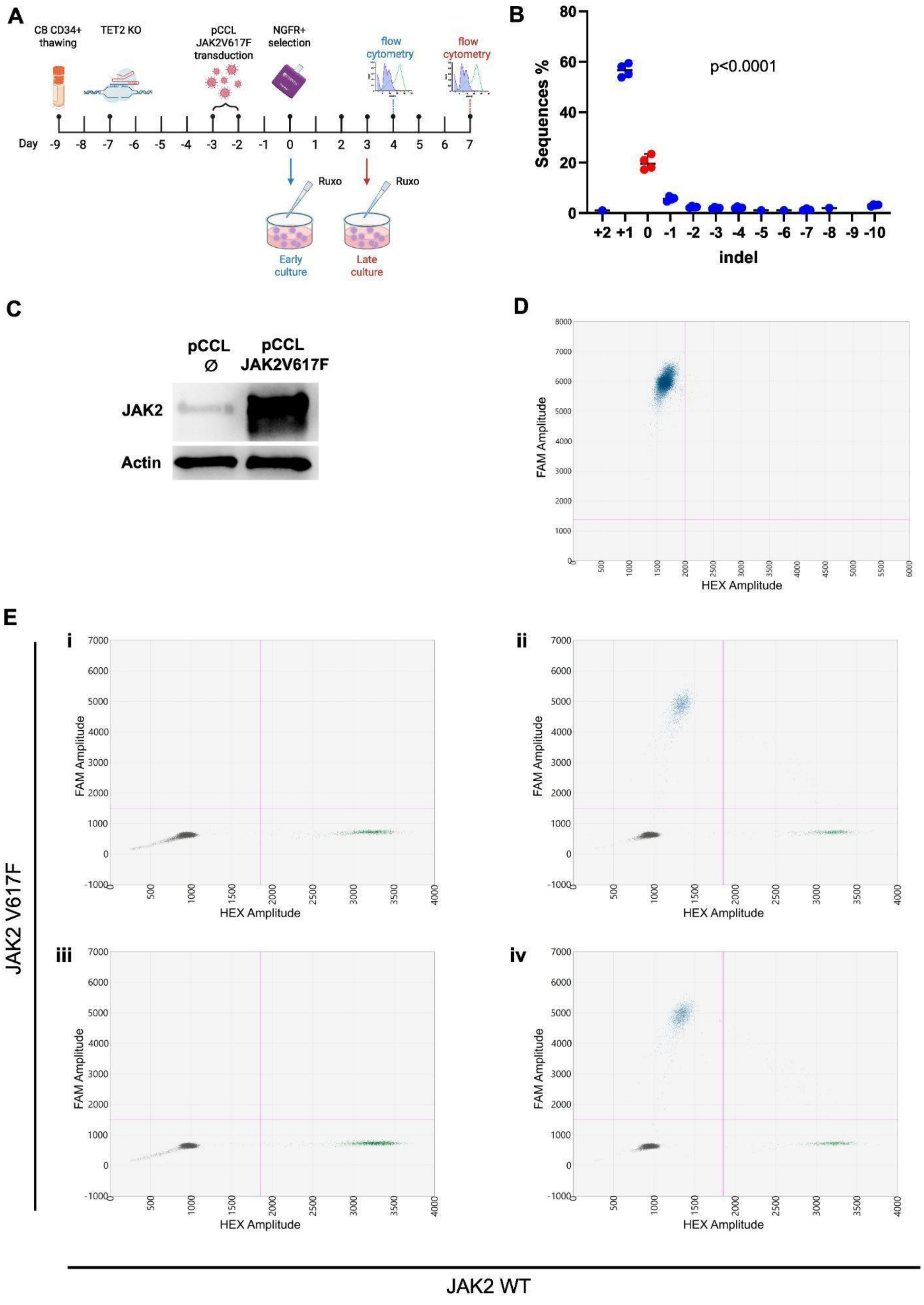


**Supplementary Figure 7**: **A** Experimental design. The same CRISPR-Cas9 strategy used for genome editing in Hel cells was used to obtain *TET2* KO in cord blood (CB) derived CD34+ cells. Cryopreserved CD34+ cells isolated from CB were thawed and 48h later the selected guide and Cas9 enzyme were nucleofected into cells. 4 days later, cells were transduced with a lentiviral vector (pCCL) to overexpress JAK2V617F MPN driver mutation. After infection, NGFR+ transduced cells were immunomagnetically sorted and seeded in multiwell plates. Purified cells were then treated daily with 2.5µM Ruxolitinib (RUXO) to evaluate the effect of JAK inhibition on growth of CD34+ cells in early culture. After 96 hours, the frequency of NGFR+ transduced cells was evaluated by means of cytofluorimetric analysis within CD34+ hematopoietic stem and progenitor cells.

In late culture experiment, 3 days after NGFR+ cells purification, cells were treated daily with RUXO and CD14+ monocytes growth was evaluated by flow cytometry after 96 hours. **B** *TET2* targeting efficiency in CB derived CD34+ cells using Guide 3. Scatter plot showing the frequency of the different in-dels generated and unmodified alleles. Each dot represents an experiment (n=4). Comparison was performed using ANOVA test. **C** Western blot evaluating JAK2 protein expression in HEK 293T cells after pCCL JAK2V617F transfection. **D** Droplet digital PCR (ddPCR) results highlighting the expression of JAK2V617F mRNA in HEK 293T transfected cells. **E** ddPCR was used to validate the expression of JAK2V617F mRNA in transduced CB derived CD34+ cells. A representative experiment is shown. Each panel displays results for a specific sample. Panels **i** and **ii** include samples nucleofected with non targeting (NT) guide, while panels **iii** and **iv** include samples nucleofected with *TET2* targeting (TET2 KO) guide. Panels **i** and **iii** include samples transduced with empty vector (pCCL Ø), while panels **ii** and **iv** include samples transduced with lentiviral vector containing JAK2V617F variant (JAK2V617F). Abbreviations: pCCLØ: sample transduced with empty pCCL lentiviral vector; pCCL JAK2V617F: sample transduced with pCCL JAK2V617F lentiviral vector; ΔNGFR+: cells expressing Truncated Nerve Growth Factor Receptor; RUXO:Ruxolitinb;

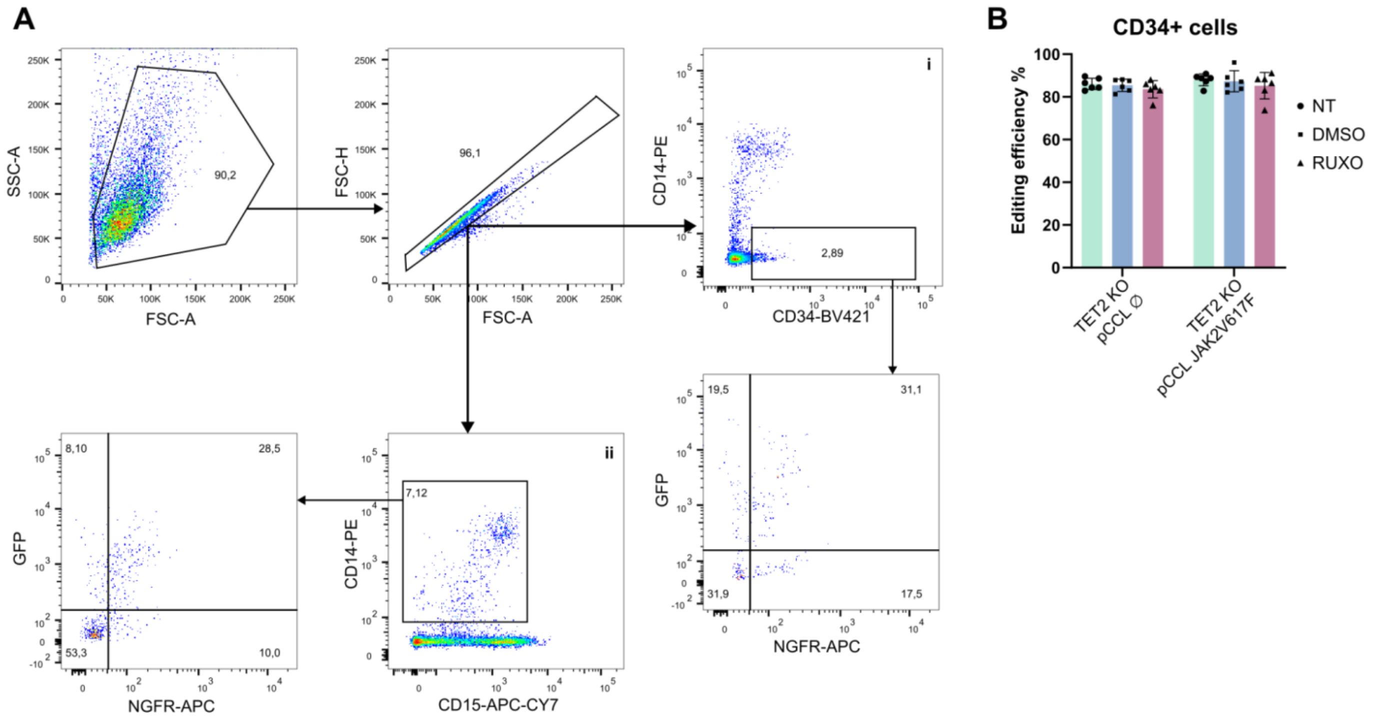


**Supplementary Figure 8** : **A** Representative flow cytometry gating strategy of transduced (ΔNGFR+) CD34 and CD14 positive cells. Cells are gated on forward (FSC) versus side scatter (SSC) and next on FSC-Area (FSC-A) vs FSC-Heigh (FSC-H) to exclude doublets. The percentage of ΔNGFR+ and/or GFP+ is analyzed among each cell population. Panel **i** shows CD34 positive cells; Panel **ii** shows CD14 positive cells. **B** Bar plot showing the editing efficiency in CD34+ cells after Ruxolitinib (RUXO) treatment. Abbreviations: DMSO: Dimethyl Sulfoxide; FSC-A: forward scatter-Area; SSC-A: side scatter-Area; SSC-H: side scatter-Heigh; TET2 KO pCCLØ: sample nucleofected with CRISPR-Cas9 *TET2*-targeting guide and transduced with empty pCCL lentiviral vector; TET2 KO pCCL JAK2V617F: sample nucleofected with CRISPR-Cas9 *TET2*-targeting guide and transduced with pCCL JAK2V617F lentiviral vector.
